# Novel regulators of nitric oxide signaling triggered by host perception in a plant pathogen

**DOI:** 10.1101/779173

**Authors:** Yi Ding, Donald M. Gardiner, Di Xiao, Kemal Kazan

**Affiliations:** Agriculture and Food, Commonwealth Scientific and Industrial Research Organization, 306 Carmody Road, St Lucia, 4067, Queensland, Australia; The School of Chemistry and Molecular Biosciences, The University of Queensland, St Lucia 4072, Queensland, Australia

**Author notes:** **Corresponding authors** Donald M. Gardiner, Kemal Kazan.

## Abstract

The rhizosphere interaction between plant roots or pathogenic microbes is initiated by mutual exchange of signals. However, how soil pathogens sense host signals is largely unknown. Here, we studied early molecular events associated with host recognition in *Fusarium graminearum*, an economically important fungal pathogen that can infect both roots and heads of cereal crops. We found that host-sensing prior to physical contact with plant roots radically alters the transcriptome and triggers nitric oxide (NO) production in *F. graminearum*. We identified an ankyrin-repeat domain containing protein (FgANK1) required for host-mediated NO production and virulence in *F. graminearum*. In the absence of host plant, FgANK1 resides in the cytoplasm. In response to host signals, FgANK1 translocates to the nucleus and interacts with a zinc finger transcription factor (FgZC1), also required for NO production and virulence in *F. graminearum*. Our results reveal new mechanistic insights into host-recognition strategies employed by soil pathogens.

## Introduction

The rhizosphere, the soil region where plant roots and a diverse array of soil microbes known as the root microbiome interact, harbors the largest reservoir of microbial diversity on earth ^1^. The majority of plant-microbe interactions in the rhizosphere is either beneficial or commensal. Successful initiation of such interactions requires exchange of molecular signals between hosts and microbes even in the absence of any physical contact between the interacting partners. For instance, beneficial interactions of plant roots with bacterial (e.g. rhizobia) ^2^ or fungal (e.g. arbuscular mycorrhiza) ^3^ organisms start with the arrival of biologically active compounds such as flavonoids, lipochitooligosaccharides and strigolactones secreted either by plant roots or microorganisms ^4^.

The rhizosphere also contains many pathogenic microbes that can pose a significant threat to plant productivity and ecosystem health. However, in a complex environment like the rhizosphere, successful colonization of plant roots by soil pathogens can be dependent on successful sensing of the host plant. Indeed, pathogenic fungi can sense the proximity of plant roots by detecting the presence of various compounds found in plant root exudates. Host-sensing triggers a developmental response known as chemotaxis that can direct the movement of the pathogen towards plant roots. A recent example of this phenomenon is the sensing of peroxidases released by wounded tomato roots by the soil-dwelling fungal pathogen *Fusarium oxysporum* ^5^. Peroxidase chemosensing requires the synthesis of reactive oxygen species (ROS) by the NADPH oxidase B complex, the G-protein coupled receptor Ste2 and the mitogen-activated protein kinase Mpk1 in *F. oxysporum* ^6^. Similarly, after sensing diverse molecules found in root exudates, bacterial pathogens *Ralstonia solanacearum* and *Agrobacterium tumefaciens* swim towards host roots using their flagella ^7–9^.

These examples clearly demonstrate that belowground signal exchanges between plant roots and soil microbes are important for successful interactions. Thus, better understanding of these processes can lead to the design of new strategies to promote beneficial interactions or combat plant diseases. However, in contrast to root-beneficial microbe interactions, very little is known about host-perception and signal exchange processes involved in plant-soil pathogen interactions ^10^. It is conceivable, nevertheless, that root-derived signals activate pathogenesis/virulence- and developmental processes in soil pathogens. Therefore, interfering with these signal exchange processes such as modifying plant-derived signals to subvert host recognition could make plant roots “invisible” to soil pathogens.

To improve our understanding of signal exchanges between plant roots and pathogenic fungi, in this study, we investigated early molecular events triggered by host-sensing in *F. graminearum* (*Fg*) using the model *Fg*-*Brachypodium distachyon* (*Bd*) interaction. The pathogenic fungus *Fg* causes some of the most economically important diseases of cereal crops and mycotoxin contaminations in food and feed products, resulting in billions of dollars of yield losses worldwide and threaten our food supply and safety ^11^. Although *Fg* is better known for its devastating effect on wheat and barley heads as well as maize stalk, recent studies have demonstrated the ability of this pathogen to act as a soil pathogen on wheat, barley, maize, soybean as well as the model monocot *Bd* ^12–17^. To study host-sensing in *Fg*, we conducted an RNA-seq analysis to identify candidate fungal genes differentially regulated by host-derived signals prior to physical contact with host roots. Functional analyses through gene-knock out analyses showed that pre-contact *Fg* genes are involved in fungal development, metabolism as well as virulence. In addition, we found that sensing of host signals triggers nitric oxide (NO) production in the pathogen. We identified two novel regulators of NO production in *Fg*, an ankyrin repeat containing protein (FgANK1) and a zinc finger transcription factor (FgZC1). We showed that these two proteins physically interact, and both are required for NO production and pathogen virulence. In the absence of host signals, no interaction between FgANK1 and FgZC1 and no NO production could be observed. Overall, our results reveal mechanistic new insights into previously unknown components of host-sensing apparatus in an important plant pathogen.

## Results

### Sensing of host signals reprograms the *Fg* transcriptome

To gain new insights into fungal processes potentially involved in the recognition of host-specific signals prior to physical contact with roots, we performed transcriptome (RNA-Seq) analyses on the interaction between *Brachypodium distachyon* (*Bd*) and *F. graminearum* (*Fg*) (Fig. 1, Supplemental data 1). These analyses focused on three stages of *Fg* growth on minimal medium (MM) designated as (i) *Fg*-only (no *Bd*), (ii) Pre-contact (*Fg* grown in the presence of *Bd* but without any physical contact with the roots) and (iii) Colonization (*Fg* infecting *Bd* roots) (Fig. 1a). Principal component analysis (PCA) conducted on the transcriptome data revealed distinct differences between fungal transcriptomes from each stage (Fig. 1b). We identified 678 (|logFC| ≥ 1, FDR < 0.05) differentially expressed fungal genes (DEGs) in the pre-contact stage. Of these, 279 genes were up-regulated in the pre-contact stage relative to the *Fg*-only stage (Fig. 1c, Supplemental data 2). Interestingly, nearly half (320) of the pre-contact DEGs were not differentially regulated at the colonization stage comparing to *Fg*-only (Fig. 1c, Supplemental data 2), suggesting that different fungal processes might be operational at different stages. The pre-contact DEGs can be grouped into six distinct clusters, which clearly distinguished the stage dependent transcription patterns (Fig. 1d).

**Figure 1.**
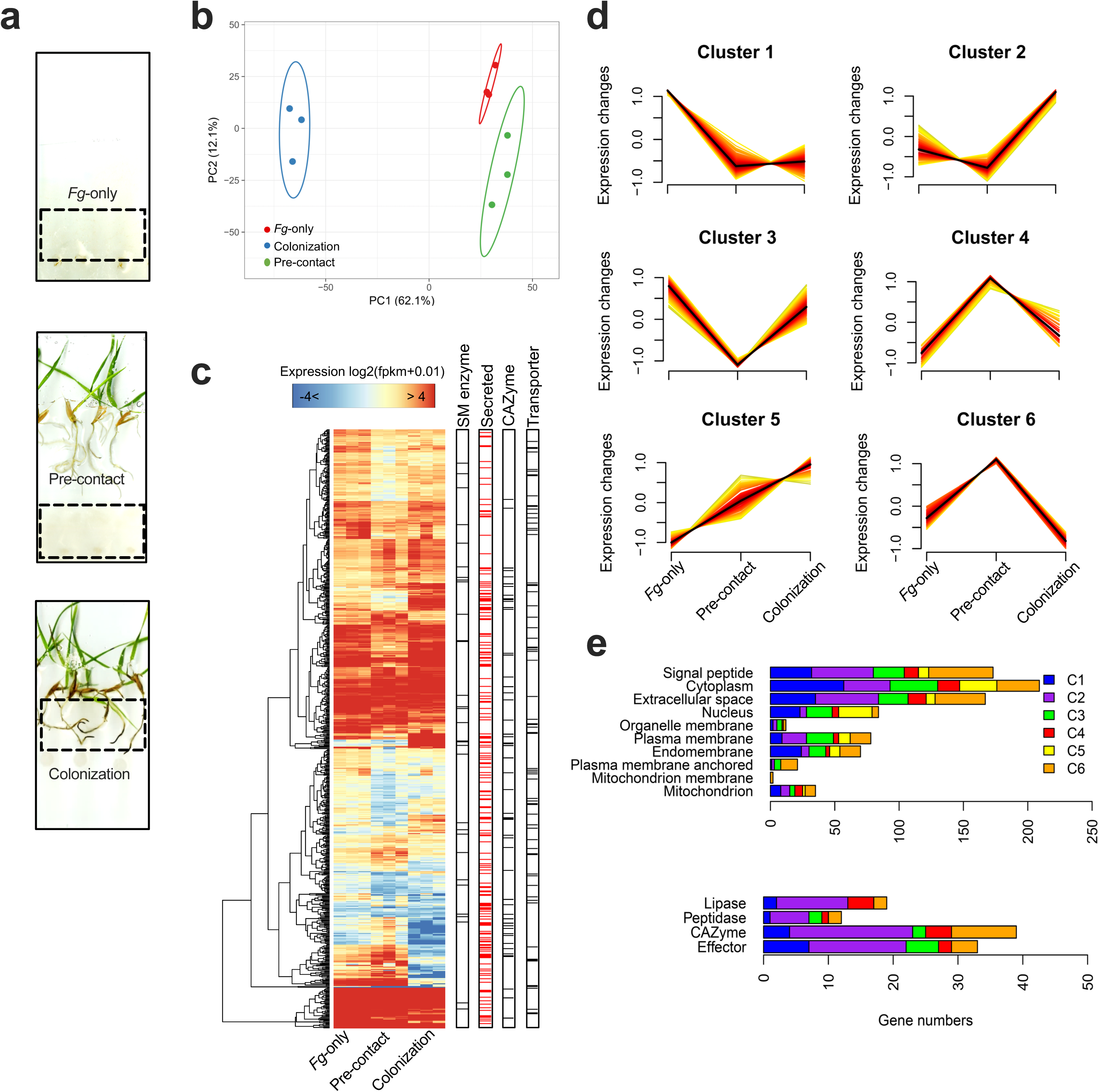
Transcriptomic reprogramming in *F. graminearum* (*Fg*) prior to contact with *Brachypodium distachyon* roots (*Bd*). *Fg* was either grown alone on minimal medium (MM) (*Fg*-only) or in the presence of *Bd* but either with or without (Colonization and Pre-contact stages, respectively) physical contact with *Bd* roots. At 7 dpi, RNA was isolated from fungal mycelia of all three stages of *Fg* growth and subjected to RNA-seq analysis. **a** The experimental system (*Fg*-only, pre-contact and colonization) used in transcriptome analyses. **b** Principal component analysis (PCA) of normalized RNA-Seq data. FPKM (Fragments Per Kilobase of transcript per Million mapped reads) values from all expressed genes were logarithmically transformed and used from *Fg-*only, pre-contact and colonization samples, each consisting of three biological replicates. X and Y axes show Principal Component 1 (PC1) and Principal Component 2 (PC2) that explained 62.1% and 12.1% of the overall variance, respectively. It is estimated that a new observation from the same group would fall inside a prediction ellipse with a probability value of 0.95. **c** Expression patterns of 678 differentially expressed genes (DEGs) (Pre-contact *vs Fg*-only, |log_2_FC|>1, FDR<0.05). A pseudo-count of 0.01 was added to the raw FPKM values for log2-transformation. Dendrograms represent distance dissimilarity across-gene expression levels and samples. Sidebars highlight key features associated with common fungal metabolic and virulence pathways. **d** Fuzzy clustering of all genes presented in **b** based on possible expression trend across the tested fungal growth stages. Mean values of FPKM were standardized to have a mean value of 0 and a standard deviation of 1, allowing similar changes in expression. **e** The number of genes from different functional categories (upper panel) or involved in putative secreted pathways (lower panel) present in all six clusters.

Transcriptome analyses suggested that *Fg* senses host-derived signals prior to physical contact with host roots and radically alters its transcriptome in anticipation of an infection. Presumably, host-derived signals altering the *Fg* transcriptome at the pre-contact stage include molecules released by *Bd*. To evaluate this hypothesis, we treated *Fg* hyphae with root exudates isolated from *Bd* plants grown in the presence or absence of the fungus and analyzed the expression of selected *Fg* genes by RT-qPCR. Interestingly, the root exudates from *Bd* seedlings grown in the presence of *Fg* altered the expression of selected pathogen genes (Fig. 2). This is consistent with the idea that the host plant senses the pathogen and alters the composition of its root exudates, which can then serve as host-recognition signals for *Fg*.

**Figure 2.**
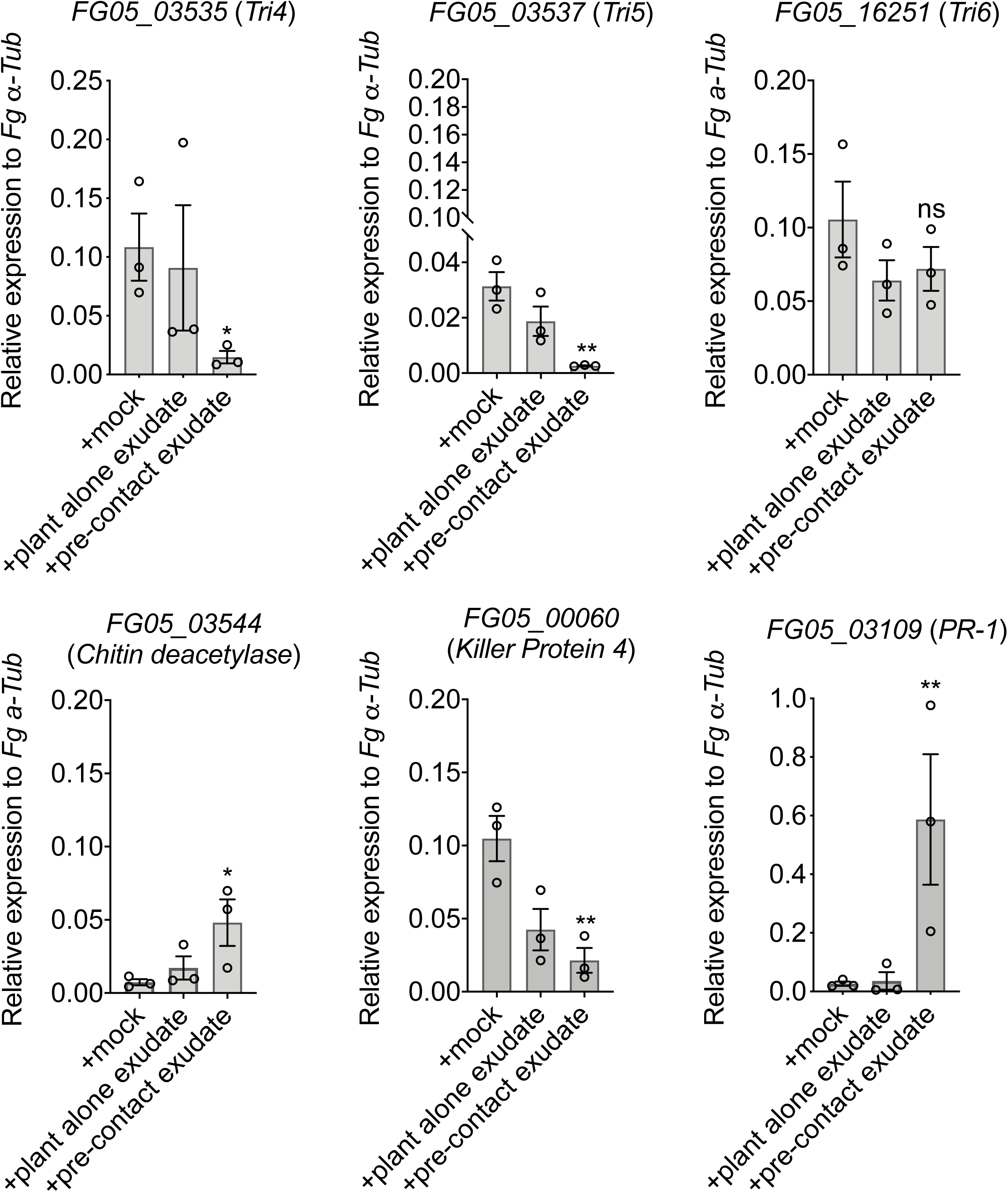
*Fg* senses host signals from *Bd* plants at the pre-contact stage (i.e. *Bd* plants grown in the presence of *Fg*) but not the *Bd* alone (i.e. *Bd* plants grown in the absence of *Fg*) stage. *Fg* grown in liquid MM was either mock-treated or treated with *Bd* root exudates isolated from plants grown in the presence or absence of *Fg*. Expressions of selected *Fg* genes were analyzed using RT-qPCR. Gene expression levels were normalized using the fungal reference gene *α-Tubulin*. Data are the mean of three biological replicates with error bars representing standard error of the mean. Asterisks represent differences that were statistically significant compared with the mock control (unpaired two-tailed *t*-test, *P< 0.05. **P< 0.01).

### Developmental processes are altered during the pre-contact stage in *Fg*

Given their potential roles of sensing environmental stimuli, pre-contact DEGs associated with pathways related to cell surface construction and signal transduction, or processes involving receptor kinases and transcription factors (TFs) were of particular interest ^4,5^. Comparing to published datasets, we identified several pre-contact DEGs involved in fungal morphogenesis and development ^18–23^. These included cell wall biosynthesis genes (*FG05_06888* and *FG05_07891* from Clusters 4 and 5), fourteen genes encoding cell surface/membrane-anchored proteins from Cluster 6 (Supplemental data 3, Fig. 1e), and putative protein kinase genes (*FG05_08701* and *FG05_02488* from Cluster 6). In addition, two upregulated TFs (FG05_11850 and FG05_03597, Supplemental data 4) were implicated in fungal conidiation and morphogenesis ^23,24^. Half of predicted *Fg* genes encoding cell wall integrity and stress response component (WSC) proteins implicated with functions in environmental sensing and morphogenesis in eukaryotic organisms ^25^ were identified from Cluster 6 (Supplemental data 5). Taken together, processes of fungal morphogenesis and development appear to be specifically regulated during the pre-contact stage in *Fg*.

### A substantial portion of pre-contact DEGs encode secreted proteins

We found that proteins encoded by 173 genes (25.5% of all pre-contact DEGs) contain signal peptides (Fig. 1e, Supplemental data 3 and 6). More than 50% of these genes were found in Cluster 2 and 6 (Fig. 1e, Supplemental data 3). In particular, Cluster 2 contained putative virulence-related genes, such as putative effectors, carbohydrate-active enzymes (CAzymes), peptidases and lipases (Fig. 1e, Supplemental data 3). In Cluster 6, we identified two highly induced genes (*FG05_00009* and *FG05_11513*) encoding putative effectors (Supplemental data 2, 8 and Supplementary Table 1). We generated independent knockout mutants for these putative effectors and tested their stress-related growth- and pathogenesis-phenotypes on *Bd* roots and wheat heads (Supplementary Fig. 1). Interestingly, while resembling to WT growth phenotypes in the absence of stress, both *FG05_00009* and *FG05_11513* mutants displayed increased sensitivity to salt stress (Supplementary Fig. 1a). *FG05_00009* mutants also showed significantly reduced virulence on wheat heads (Supplementary Fig. 1b and 1c), suggesting that pre-contact regulated genes encoding putative effectors may be involved in stress tolerance and pathogen virulence in *Fg*.

### Putative virulence and defense genes are differentially-regulated at the pre-contact stage in *Fg*

Cluster 3 was enriched in fungal virulence, which included three top down-regulated pre-contact genes encoding secreted fungal killer protein (KP4) (Supplemental data 7 and 8) shown to be involved in virulence against wheat ^26^. In addition, *FG05_04745* encoding an antifungal-related effector was strongly-downregulated in Cluster 2 (Supplemental data 8). The induction of a predicted chitin deacetylase encoding gene (*FG05_03544*) from Cluster 4 and a plant pathogenesis-related (PR-1) protein homolog gene (*FG05_03109*) from Cluster 6 (Supplemental data 8) were also observed. Fungal chitin deacetylases and PR-1 proteins may play important roles in self-protection against hydrolases ^27,28^. Surprisingly, the trichothecene biosynthesis genes *Tri4* and *Tri5* involved in the production of the trichothecene toxin deoxynivalenol (DON) were suppressed pre-contact with the host (Cluster 2, Fig. 2, Supplemental data 2 and 9).

### Nutrient metabolism is altered during the pre-contact stage in *Fg*

Several pre-contact DEGs involved in glyoxylate cycle and acetyl-CoA metabolic processes were identified in Cluster 4 (Supplemental data 7), suggesting that energy remobilization processes from fatty acids are operational at this stage. Many of these genes were previously shown to be involved in asexual development in *Fg* ^29,30^. In addition, a large number of significantly downregulated transporter-encoding genes (49 out of 71) mostly associated with sugar and amino acid/polyamine transport were found in Clusters 2 and 3 (Supplemental data 10). Cluster 2 was also enriched for genes involved in carbohydrate, tryptophan and GABA metabolic processes (Supplemental data 7), suggesting that drastic alterations in nutrient metabolism occur at this stage.

In Clusters 4 and 6, we found pre-contact DEGs enriched for nitrate, ammonium and urea transport (Supplemental data 7 and 10). *Fg* contains only two genes (*FG05_03111* and *FG05_08884*) encoding putative urea permeases (DURs), which mediate urea and polyamines transport in many Ascomycota fungi ^31–34^. In *Candida* spp., DUR31 was shown to function as a novel transporter of thiamine ^35^, an essential co-factor for metabolic and energy pathway enzymes in all living organisms ^36^. FG05_03111 and FG05_08884 show 74% and 23% identity to *C. albicans* DUR3 and DUR31, respectively, and thus were named as FgDUR3 and FgDUR31. Individual knockouts for *FgDUR3* and *FgDUR31* were generated and tested for growth on MM supplemented with glucose and different N sources. None of the single deletions affected fungal growth when urea or spermidine was used as the sole N source (Supplementary Fig. 2a). In fact, N source is not essential for *Fg* growth as indicated by normal growth of fungal strains on MM without N supplementation (Supplementary Fig. 2a). However, the growth of the FgDUR31 mutant was significantly inhibited in the presence of 1mM NH_4_NO_3_, suggesting a potential role of FgDUR31 in N responses. *FgDUR31* deletion also allowed fungal growth in the presence of the toxic thiamine analog pyrithiamine (Supplementary Fig. 2a), suggesting that FgDUR31 is involved in thiamine transport. While no growth inhibition could be detected in either mutant under various stress treatments (Supplementary Fig. 2b), only the mutant with *FgDUR31* deletion showed reduced pathogenicity towards wheat heads and *Bd* roots (Supplementary Fig. 2c). In *C. albicans*, DUR31 can mediate environmental alkalization and hyphal morphogenesis during multiple infection stages despite its downregulation ^33^. FgDUR31 might possess a similar regulation pattern affected by environmental cues and involved in disease development. Indeed, the closely related pathogen *F. culmorum* induces alkanization of the wheat host environment early during infection ^37^.

### NO biosynthesis is activated during the pre-contact stage in *Fg*

In addition to N assimilation, genes associated with nitric oxide (NO) biosynthesis and response to nitrosative stress were significantly enriched among pre-contact DEGs (Supplemental data 7). These genes include a nitrate reductase (*NR*) (*FG05_01947*) and a flavohaemoglobin NO dioxygenase (*FHB*) (*FG05_00765*) potentially involved in NO production and metabolism, respectively ^38^. In the model fungus *Aspergillus nidulans*, NR homologs are closely associated with NO homeostasis and co-regulated with nitrate assimilatory genes functionally independent of N metabolite repression ^38,39^. Moreover, intra- and extra-cellular NO directly affect the expression of conidiogenesis-related genes and fine-tune hyphae and conidia development in many fungal species ^38,40–42^. Consistent with this, the conidiation regulator ortholog BrlA (FG05_01576) from Cluster 4 was co-regulated with the NO-associated and N transporter genes (Supplemental data 4).

To determine if NO is produced in *Fg* during the pre-contact stage, hyphae from *Fg*-alone and pre-contact stages were stained using 4-amino-5-methylamino-2,7-difluorofluorescein diacetate (DAF-FMDA), which detects NO ^43,44^. To ensure that the staining by DAF-FMDA was specific for NO, DAF-FMDA with or without the cell-permeable NO scavenger 2-(4-car-boxyphenyl)-4,4,5,5-tetramethylimidazoline-1-oxyl-3-oxide (cPTIO) ^42^ was applied to the edge of the mycelium growing towards the roots or to the corresponding locations in the *Fg*-alone plate. DAF-FMDA staining revealed predominant fluorescence signals in hyphal tips and branches of *Fg* during the pre-contact stage (Fig. 3b). In contrast, no such signal could be detected in fungal-alone or cPTIO treated samples (Fig. 3a and 3c). Furthermore, NO production was also triggered when the pre-contact *Bd* root extract was supplemented to the fungus grown in liquid MM (Fig. 3f and 3h). As expected, no NO production was evident in *Fg* grown either in the absence of pre-contact *Bd* root exudates or in the presence of plant signals and cPTIO (Fig. 3d-e, 3g-h). Thus, endogenous NO production in *Fg* during the pre-contact stage is triggered by host signals.

**Figure 3.**
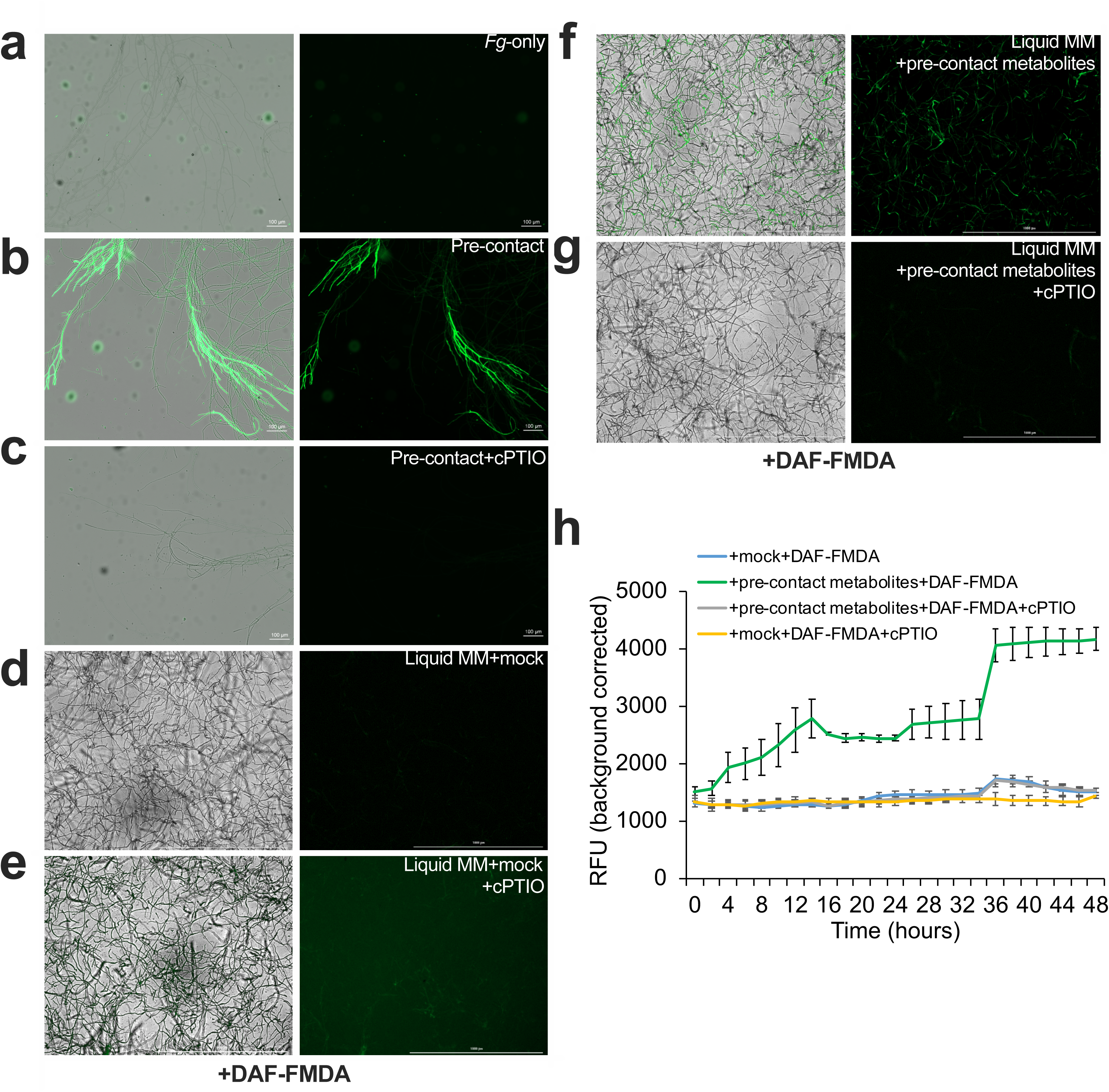
Nitric oxide (NO) is produced in *Fg* in response to signals derived from *Bd* roots. **a-b** 4-amino-5-methylamino-2,7-difluorofluorescein diacetate (DAF-FMDA) was applied to the edge of the *Fg* mycelium growing either alone (**a**) or towards *Bd* roots (**b**). DAF-FMDA-stained mycelia were rinsed briefly in water and visualized under a fluorescence microscopy for NO production. No NO-specific fluorescent signals could be observed when the fungus was grown alone (**a**) while fluorescent signals were predominantly detected in hyphal tips prior to physical contact with *Bd* roots (**b**). **c** NO-scavenger 2-(4-carboxyphenyl)-4,4,5,5-tetramethylimidazoline-1-oxyl-3-oxide (cPTIO) inhibited NO accumulation observed at the pre-contact stage. **d-g** *Bd* root exudates trigger NO biosynthesis in *Fg*. Ten µl of 10^6^ /ml of *Fg* spores were collected and transferred into 200 µl of liquid MM and analyzed by a fluorescence plate reader. NO was detected when *Fg* was treated with pre-contact metabolite extracts (**f**), but not with mock (**d** and **e**) or cPTIO treatments (**e** and **g**). **h** Kinetic analysis of NO production by *Bd* root exudates. *Fg* grown in liquid MM in the presence or absence of *Bd* root extracts were treated with DAF-FMDA and/or cPTIO and NO production was assayed over a 48-h time-course using a fluorescence plate reader. Data are the mean of three biological replicates with error bars representing standard error of the mean.

Previous studies in several fungal species have revealed roles for NO in global control of transcriptional and metabolic reprogramming and in modulating fungal morphogenesis and secondary metabolism ^45–48^. To test if NO is involved in transcriptional regulation in *Fg* during the pre-contact stage, the NO inducibility of *NR* as well as several other pre-contact DEGs was analyzed in *Fg* by RT-qPCR after the application of the NO chemical donor S-Nitroso-N-acetyl-DL-penicillamine (SNAP). Indeed, SNAP treatment mimicked the effect of the pre-contact treatment on the regulation of the selected genes (*Tri5*, *FHB*, *FG05_00416*, *FG05_11858* and *FG05_00060*), which could be compromised by cPTIO (Supplementary Fig. 3). In contrast, SNAP treatment did not alter *NR*, while the *FHB* gene potentially involved in NO detoxification ^38^ was responsive to SNAP (Supplementary Fig. 3). Among the genes potentially involved in NO biosynthesis, *NR* (*FG05_01947*) is the only possible mediator significantly up-regulated during the pre-contact stage, indicating that NR might be associated with NO production in *Fg*. The lack of induction by NO may suggest that NR is upstream of the NO signal and that no positive feedback exists if NR is indeed the source of NO in *Fg*. Nevertheless, the regulation of the remaining marker genes by SNAP suggests that NO may at least partially contribute to the transcriptional responses observed during the pre-contact stage in *Fg*.

### An ankyrin-repeat containing protein regulates NO biosynthesis during the pre-contact stage in *Fg*

The findings presented above suggests that NO is produced in *Fg* in response to host signals at the pre-contact stage although potential regulators acting upstream of NO production are unknown. Therefore, we next focused on the identification of regulatory genes that may be involved in host-sensing mediated NO production in *Fg*. We selected 19 candidates belonging to the ‘binding’, ‘signal’ and ‘unknown’ functional categories among the top differentially regulated genes at the pre-contact stage and generated their independent deletion mutants. Of these, 16 mutants exhibited at least one phenotype of altered mycelium growth, stress tolerance or virulence, while 15 showed considerably reduced virulence in root infection assays. Seven of these mutants also showed defects during wheat head infections (Supplementary Table 1, Supplementary Fig. 4), suggesting that shared pathogenicity processes might be used during the infection of different host tissues by *Fg*. Notably, we found that NO production observed at the pre-contact stage was completely abolished in the *FG05_02877* (named here as *FgANK1*) mutant (Supplementary Fig. 5c), but not the other deletion mutants (Supplementary Table 1). Besides, ΔFgANK1 was no longer responsive to *Bd* root exudates as evidenced by unchanged expression of the fungal marker genes (*Tri4*, *Tri5*, *FHB*, *FG05_03544*, *PR-1* and *FG05_00060*) (Supplementary Fig. 6a). We noticed that the deletion of *FgANK1* also resulted in abnormal pigmentation and impaired growth on all tested stress conditions (Fig. 4a) but the growth rate of this mutant on PDA was not altered (Supplementary Fig. 5a). ΔFgANK1 also exhibited an aberrant hyphal morphology with significant apical hyperbranching (Fig. 4b). In addition, no disease symptom caused by ΔFG05_02877 on *Bd* roots or wheat heads could be observed at 10 dpi (Fig. 4c-d). Such drastic reduction in pathogen virulence was not simply due to delayed disease development as there was no obvious disease symptom development even three weeks after direct root inoculations with ΔFG05_02877 (Supplementary Fig. S5b). These phenotypes could be fully complemented by expressing *GFP-FG05_02877* fusion constructs in ΔFG05_02877 (Fig 4a, 4c-e, Supplementary Fig. 6b). Interestingly, however, the mutant was responsive to external NO as the SNAP treatment resulted in similar regulation patterns of the genes *Tri5*, *FHB*, *FG05_00416* and *FG05_00060* in ΔFG05_02877 and the complemented strain (Supplementary Fig. 6b). Together, these findings suggested that FgANK1-mediated NO production is responsible for the regulation of downstream genes although similarly to *NR*, *FG05_02877* was not regulated by external NO treatment in the WT pathogen (Supplementary Fig. 5d and 6b).

**Figure 4.**
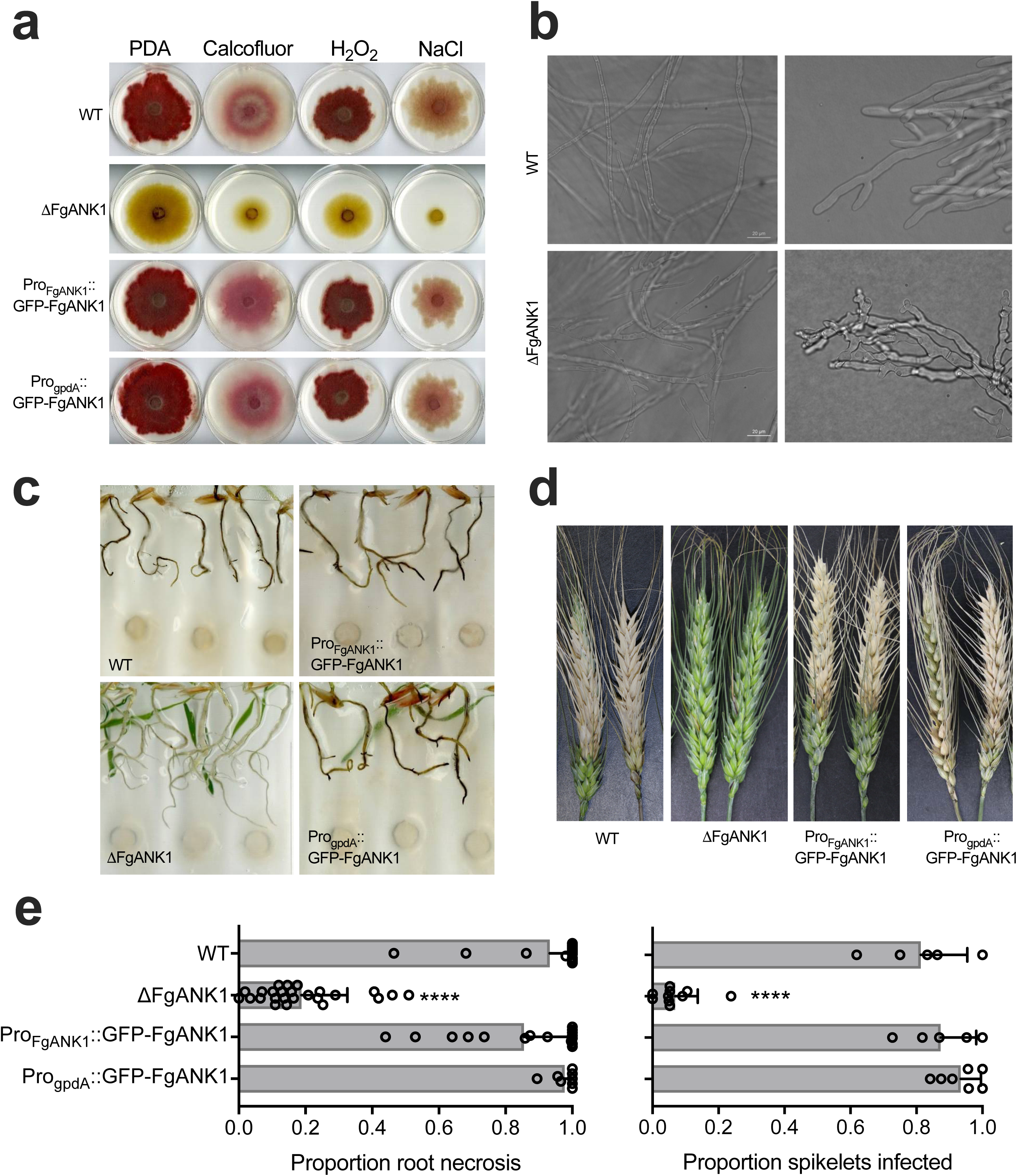
FgANK1 is required for hyphal development and virulence in *Fg*. **a** ΔFG05_02877 shows abnormal pigmentation and increased sensitivity to cell wall (0.5 mg/ml Calcofluor), oxidative (2 mM H_2_O_2_) and osmotic (0.7 M NaCl) stresses. These defects could be complemented with the expression of GFP fusion proteins GFP-FgANK1 in the mutant background using either native *FgANK1* or the *gpdA* promoter. **b** ΔFG05_02877 exhibits aberrant hyphal morphology with significant apical hyperbranching relative to WT fungus**. c-e** Deletion of *FG05_02877* significantly reduced *Fg* virulence towards *Bd* roots (**c** and **e**) or wheat heads at 10 dpi (**d** and **e**). Virulence-associated defects observed in ΔFG05_02877 could be restored by the expression of GFP fusion constructs Pro*_FgANK1_*:: *GFP*-*FgANK1* or Pro*_gpdA_*::*GFP*-*FgANK1* in the mutant background. **e** Shown here are representative data from one of at least two independent fungal transformants. Percentage *Bd* root necrosis and the number of diseased wheat spikes are averages from a minimum of twenty replicate roots and at least five wheat heads, respectively, with error bars representing the standard deviation (One-way ANOVA with Dunnett’s multiple comparisons test, ****P< 0.001).

### FgANK1 is localized in the cytosol in the absence of host-signals but translocates to the nucleus in a host-sensing mediated manner

Although the data presented above indicates that FgANK1 is a novel regulator of NO production triggered by host sensing, how FgANK1 regulates this process is not clear. FgANK1 harbors three N-terminal ankyrin-repeats and a C-terminal von-Willebrand factor type A (VWA) domain (Supplementary Fig. 7b). Ankyrin-domain containing proteins perform various functions such as transcriptional regulation, cytoskeletal organization, cellular development and differentiation, exclusively through specific protein−protein interactions mediated by the ankyrin repeat domain ^49^. Similarly, the VWA conserved domain is found in both extra- and intra-cellular proteins participating in numerous biological events and interactions with various ligands in a wide range of taxa ^50^. Indeed, FgANK1 homologs are mainly present in Ascomycota, including *Aspergillus* species and several root-associated fungi such as *F. oxysporum*, *Nectria haematococca* and *Trichoderma virens* (Supplementary Fig. 7a). No signal peptide or transmembrane domain could be found in FgANK1 (Supplementary Fig. 7b). However, we observed GFP signals exclusively localized in the cytosol when the Pro*_FgANK1_*::*GFP-FgANK1* construct was introduced into ΔFgANK1growing on MM (Fig. 5a). Interestingly, a proportion of the GFP-FgANK1 protein was translocated to the nucleus upon perception of host signals (Fig. 5b) or in the Pro*_gpdA_*::*GFP-FgANK1* strain growing *in vitro* (Fig. 5c). This subcellular localization of FgANK1 could be further confirmed by analyzing nuclear and cytosolic protein extracts (Fig. 5d). Together, these results suggest that FgANK1 mostly resides in the cytoplasm in the absence of the host plant. In response to host signals, however, FgANK1 translocates into the nucleus and regulates NO production.

**Figure 5.**
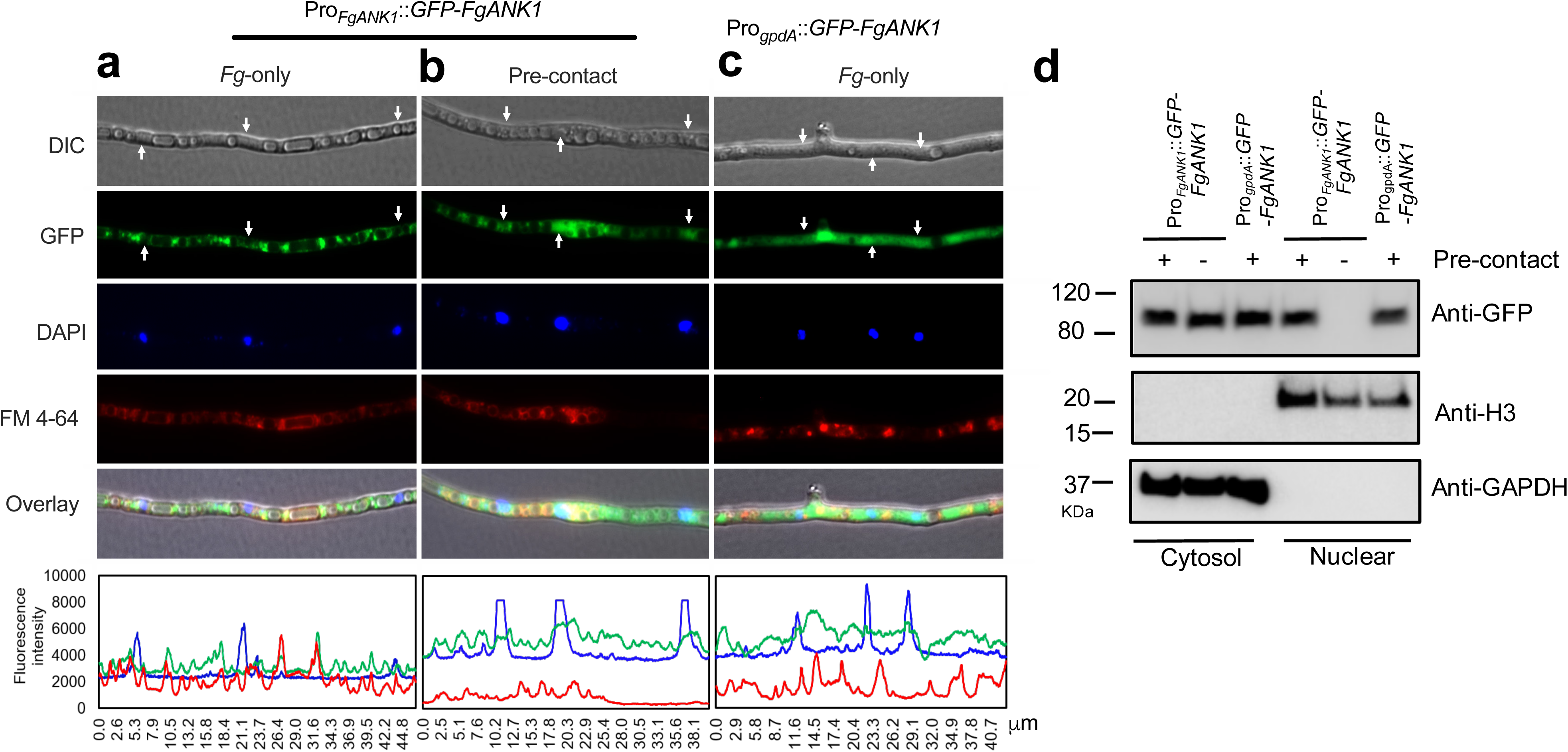
Subcellular localization of FgANK1 in *Fg*. **a**-**c** The FgANK1 ankyrin repeat domain containing protein encoded by the FG05_02877 locus accumulates in the cytoplasm during the *Fg*-only stage and both the cytoplasm and the nucleus of the *Fg* hyphae during the pre-contact stage. A FgANK1-GFP fusion protein expressed under the native *FgANK1* promoter was exclusively localized in the cytosol when the strain was grown alone on minimal medium (MM) (**a**). Cytosolic GFP signals are partially overlapped with the nuclear DAPI (4,6-diamidino-2-phenylindole) signals at the pre-contact stage (**b**) or in the overexpression strain Pro_gpdA_::GFP-FgANK1 growing in vitro (**c**). White arrows indicate nuclear positions. Fungal membranes were stained with FM 4-64. Differential Interference Contrast (DIC) images and florescence signals and intensities were shown in the top and bottom panels, respectively. **d** Western blot analyses of nuclear and cytosolic protein extracts confirm the cellular localization of FgANK1. Plus (+) and minus (-) represent the growth of fungal strains in the presence or absence of *Bd*, respectively. Cytosolic and nuclear proteins were detected using the monoclonal anti-GFP, anti-H3 and anti-GAPDH antibodies.

### The interaction between FgANK1 and a zinc finger TF regulates NO synthesis and pathogenesis in *Fg*

Since sequence analyses did not reveal any known DNA binding motifs in FgANK1, its nuclear accumulation might involve interaction with nuclear proteins. To identify a possible interaction partner of FgANK1, we screened a yeast-two hybrid (Y2H) library generated using the fungal pre-contact cDNAs. Several-rounds of high stringency screening identified a GAL4-type Zn_2_-Cys_6_ zinc-finger TF (FG05_05068 named here as FgZC1) as an interaction partner of FgANK1. We also tagged FgANK1 with blue fluorescent protein (TagBFP) and introduced into the ΔFgANK1 background for further analysis. In the absence of *Bd* roots, we found similar cytoplasm localized pattern of TagBFP-FgANK1 (Supplementary Fig. 8a), which fully complemented the phenotype of ΔFgANK1 (Supplementary Fig. 8d), indicating that TagBFP-FgANK1 functions the same as GFP-FgANK1. Further Y2H assays and co-immunoprecipitation (co-IP) using fungal transformants co-expressing FgZC1-HA and TagBFP-FgANK1 or TagBFP-FgANK1^ΔVWA^ revealed that the VWA domain of FgANK1 is not required for the interaction between FgANK1 and the zinc finger TF (Fig. 6a-b) while the ankyrin domain of FgANK1 is required for the nuclear interaction between FgANK1 and FgZC1. Importantly, the interaction between these two proteins were completely root pre-contact dependent (Fig. 6b). Interestingly, deletion of FgZC1 did not cause any defect in growth or pathogenicity towards wheat heads (Supplementary Fig. 4), but, similarly to the deletion of FgANK1, significantly affected NO production (Fig. 7a-b and 7f) and delayed *Bd* root infection (Supplementary Fig. 9a-b). During the pre-contact stage, the selected fungal marker genes were similarly mis-regulated in *FgANK1* and *FgZC1* mutants (Supplementary Fig. 9c), suggesting that both FgANK1 and FgZC1 are required for their expression at the pre-contact stage. While the expression of both TagBFP-FgANK1 or TagBFP-FgANK1^ΔVWA^ driven by the native *FgANK1* promoter in the ΔFgANK1 background complemented the defects in NO production (Fig. 7c, 7e-f) and root colonization (Supplementary Fig. 8d-e), the latter failed to complement its abnormal hyphal morphology (Fig. 7e). In contrast, the expression of TagBFP-FgANK1**^Δ^**^ANK^ fully recovered the growth phenotype but not the pre-contact NO accumulation and root pathogenicity phenotypes (Fig. 7d and 7f, Supplementary Fig. 8f). A possible explanation for this is that the disruption of the ankyrin domain region of FgANK1 resulted in a loss of interaction capability, and consequently impaired NO production. There might also be a close correlation between early NO production and fungal virulence as demonstrated by the attenuated virulence phenotypes observed in FgZC1 (Supplementary Fig. 9a-b) and FgANK1 mutants with either full-length or ANK repeats deletions of FgANK1 (Fig. 7, Supplementary Fig. 5, 8f-g). Overall, these results support a model where upon perception of host signals FgANK1 and FgZC1 physically interact to regulate NO signaling and control the transcriptional responses to host recognition (Fig. 8).

**Figure 6.**
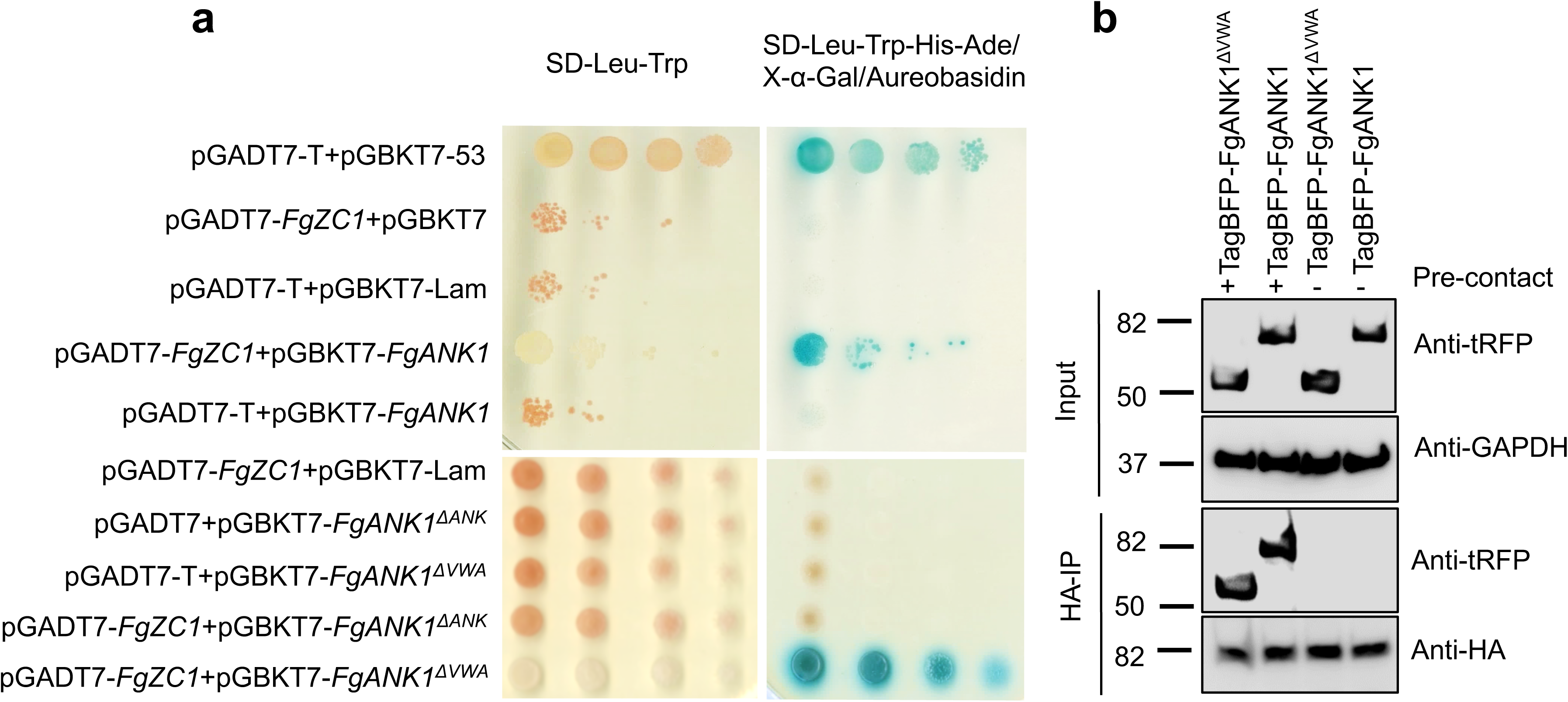
FgANK1 and FgZC1 interact in a host-sensing mediated manner in *Fg.* Y2H assays showing that the ankyrin repeats domain of FgANK1 is required for its interaction with FgZC1 at the pre-contact stage in *Fg*. **a** Y2HGold yeast cells co-expressing the bait vectors pGBKT7-*FgANK1*, pGBKT7-*FgANK1*^Δ^*^ANK^*, and pGBKT7-*FgANK1* ^Δ^*^VWA^* with the prey vector pGADT7-*FgZC1* in pairs were spotted in 10×dilutions on control (-Leu/-Trp) and on high stringency (-Leu/-Trp/-His/-Ade/+X-α-Gal/+Aureobasidin) media. The interaction of proteins expressed by the bait and prey constructs activates α-galactosidase in the presence of X-α-Gal and thus generates blue-coloured colonies. Cells expressing vector pairs pGADT7-T/pGBKT7-53 and pGADT7-T/pGBKT7-Lam were used as positive and negative controls. Self-interaction was tested by co-transforming pGBKT7-*FgANK1* or pGADT7-*FgZC1* with the corresponding control vectors. **b** Total proteins (input) extracted from *Fg* strains co-expressing FgZC1-HA with TagBFP-FgANK1 or TagBFP-FgANK1^ΔVWA^ were immunoprecipitated with the anti-HA antibody (IP). Western blots were developed with the anti-tRFP antibody to detect the interaction of FgZC1 with the full length or VWA-domain truncated FgANK1 fused with TagBFP. The FgANK1-FgZC1 interaction only occurs in the presence of *Bd* roots (pre-contact). Anti-GAPDH and anti-HA antibodies were used as loading controls for the input and IP, respectively.

**Figure 7.**
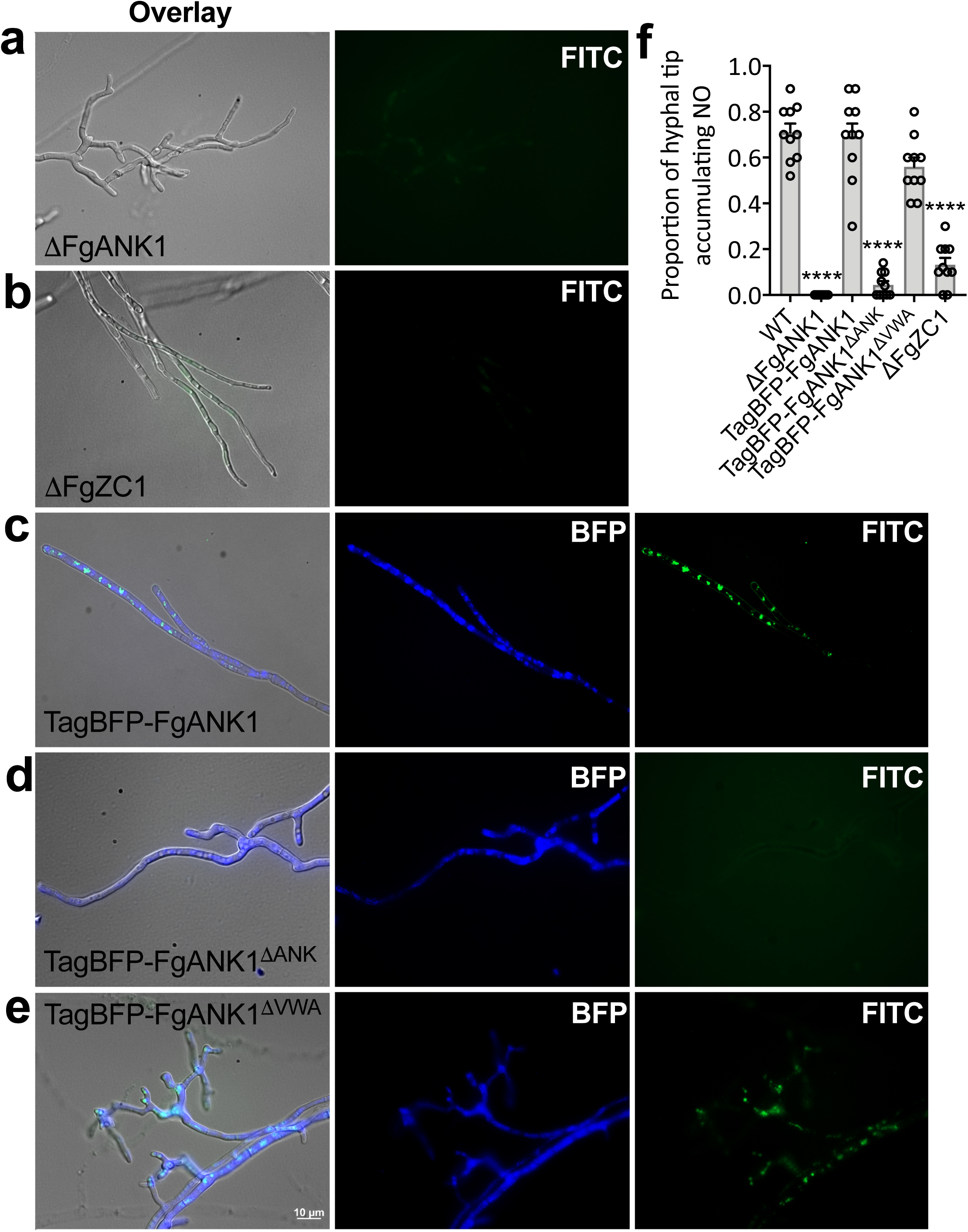
FgZC1 and FgANK1 are indispensable for NO production triggered in *Fg* by *Bd* host signals at the pre-contact stage. **a**-**c** No NO signals could be detected by FITC (Fluorescein isothiocyanate) staining in FgANK1 (**a**), and FgZC1 mutants (**b**), respectively while a *TagBFP-FgANK1* construct expressed in the ΔFgANK1 mutant background restores NO production (**c**) when *Fg* hyphae from the pre-contact stage were stained with DAF-FMDA. **d**-**f** The ankyrin repeat domain but not the VWA domain of FgANK1 is necessary for NO production at the pre-contact stage as the TagBFP-FgANK1 fusion protein with a deleted VWA region (TagBFP-FgANK1^ΔVWA^) restores NO production when expressed in ΔFgANK1 (**d**) while NO signals were completely missing in the TagBFP-FgANK1^ΔANK^ with a deleted ankyrin (ANK)-repeats domain expressed in the FgANK1 mutant background (**e**). Percentage of *Fg* hyphae accumulating NO at the apical branches of the indicated genotypes was quantified by observing ten independent microscopic slides holding stained mycelium from five inoculation plates (**f**). Data represent averages with standard error of mean (One-way ANOVA with Dunnett’s multiple comparisons test, ****P< 0.001).

**Figure 8.**
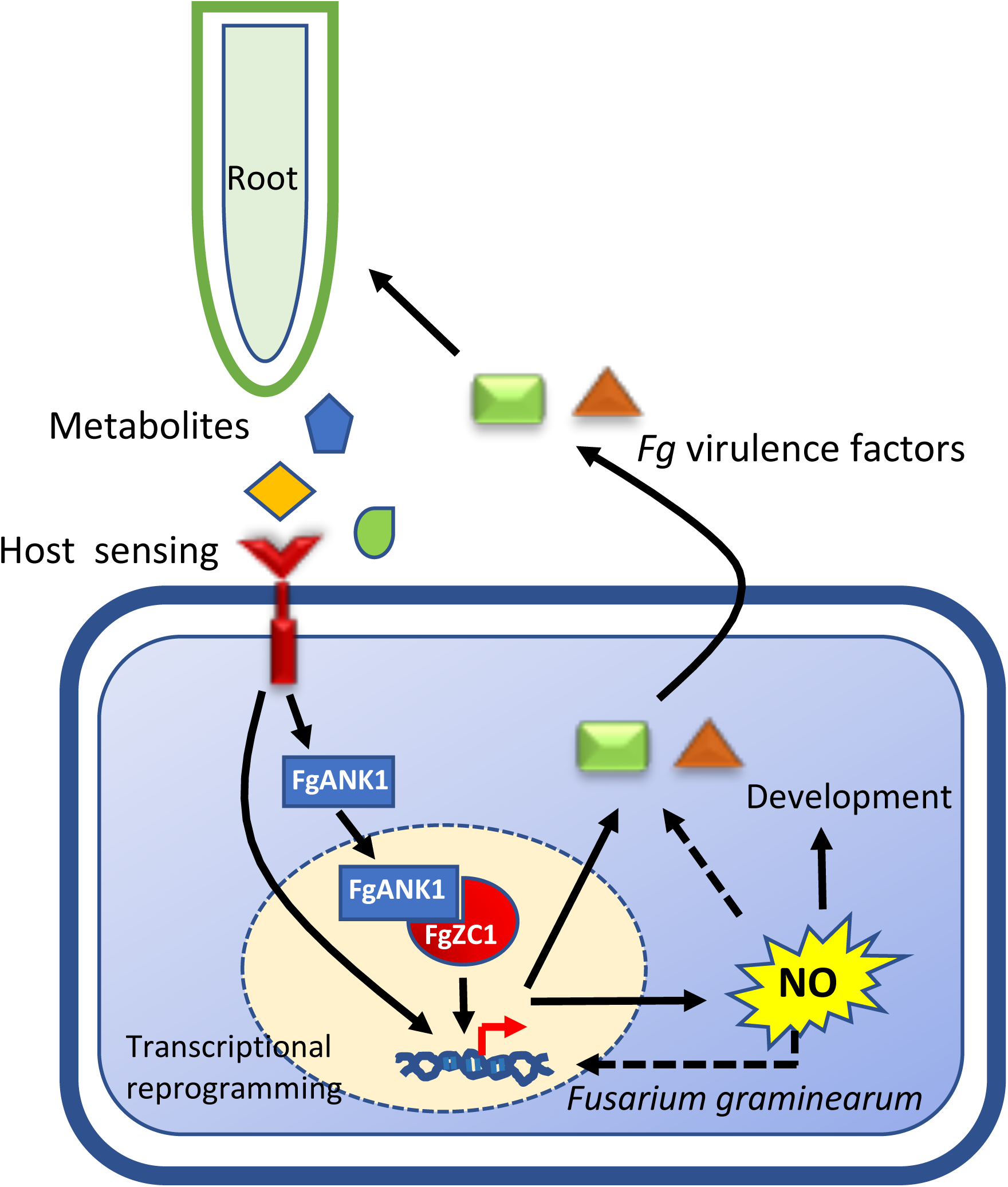
A model explaining the proposed roles of FgANK1 and FgZC1 in host-sensing mediated nitric oxide (NO) biosynthesis in *Fg*. Host-sensing triggers transcriptional re-programming and NO production in *Fg*. The ankyrin repeat domain containing protein FgANK1 resides in the cytoplasm in the absence of the host plant. Upon sensing of host signals, such as the metabolites found at root exudates, FgANK1 translocates to the nucleus and physically interacts with the zinc finger transcription factor FgZC1 to regulate NO production and pathogen virulence. NO in turn may regulate the expression of genes involved in fungal virulence and development either directly or indirectly by further modulating the transcriptome. Broken arrows refer to tenuous links.

## Discussion

During their co-evolution with plants, certain soil microbes have evolved to exploit signals released by plant roots for their benefits. Some of the best studied chemical signals associated with belowground microbe-root communications are from mutualistic interactions ^51^. Host-sensing by soil pathogens can also be critical for root infection, but the mechanisms that soil pathogens use to sense and trace host signals remain largely unexplored. Presumably, sensing of host plant in a complex environment like the rhizosphere takes places relatively early on and even in the absence of a physical interaction between the pathogen and plant roots.

In this study, we used the *Fg*-*Bd* interactions to study host-sensing mediated molecular processes in the pathogenic fungus *Fg*. To the best of our knowledge, early responses triggered by host signals in *Fg*, a fungal pathogen that can infect multiple host tissues including roots, have not been previously examined. Here, we performed detailed transcriptome analyses of different fungal growth stages and identified large number of genes differentially regulated in the presence of root signals. Our discovery on the regulation of endogenous NO during the host sensing stage sheds light on the interplay between signal recognition and developmental transitions in *Fg*, processes that are essential for both environmental adaptation and pathogenesis of this soil-borne fungal pathogen. NO-based regulatory protein modifications may mediate cellular responses involving a complex regulatory network underlying NO signaling. NO can also serve as a key messenger and a signaling molecule favoring fitness of both fungal partner and plant hosts during pathogenic as well as mutualistic interactions ^40,52–57^. Consequently, NO would deliver key messages from environmental signals and mediate the pathogen cellular responses through transcriptional reprogramming to facilitate lifestyle transition and niche adaptation ^45–48^.

Functional analyses of selected candidate pre-contact DEGs have led to the discovery of FgANK1, an ankyrin repeat domain containing protein that regulates host-sensing mediated NO production and virulence in *Fg*. Our phylogenetic analyses showed that FgANK1 homologs are found in other fungal species that include other soil-borne pathogens such as *F. oxysporum* (Supplementary Fig. 7a-b). The FgANK1 mutants were developmentally affected and incapable of infecting *Bd* roots and wheat heads. These observations suggest that FgANK1 is required for NO production as well as fungal development and virulence. It is possible that virulence and developmental alterations observed in the FgANK1 mutant could be due to defects in NO biosynthesis as NO has been implicated in these fungal processes in other fungi ^38,41,42,48,55^. It is well documented that NO is an important signaling molecule that contributes to pathogen survival within the host ^58^, and thus the loss-of-virulence phenotype of ΔFgANK1 could be due to the extreme sensitivity of this mutant to plant-defenses or the requirement of fungal-produced NO to interfere with plant defenses.

We showed that FgANK1 is a cytosolic protein and thus most likely acts downstream of a membrane receptor involved in direct sensing of host metabolites (Fig. 8). Sensing of host signals by specific receptors may activate a phosphorylation cascade that in turn can mediate the translocation of FgANK1 into the nucleus although we cannot rule out the possibility that a host metabolite taken up by the fungus directly binds to the VWA domain known to function as a ligand binding domain. Alternatively, FgANK1 translocation into nucleus may be mediated by NO-mediated redox changes. In addition, we identified the Zn2-Cys6 zinc finger TF FgZC1 required for NO production and fungal virulence as an interaction partner of FgANK1 in the nucleus. We show that the ankyrin repeat domain of FgANK1 is required for the interaction with FgZC1. Interestingly, the interaction between these proteins occurs only in the presence of host signals. FgZC1 homologs can also be found in a large range of fungal species, but clearly fall into two distinct phylogenetic clusters that comprise fungal pathogens and symbionts (Supplementary Fig. 7c).

Overall, our results suggest that the FgANK1-FgZC1 protein complex most likely acts as a master regulatory complex contributing to NO production, thus could be central for the downstream transcriptional programs (Fig 8). Although we show that NO is produced in *Fg* in a host-recognition dependent manner, and this is regulated by FgANK1 and FgZC1, the exact biosynthetic pathway by which NO is produced in *Fg* and other fungi is not clear. In *A. nidulans*, NR seems to catalyze NO production whose induction is in a nitrogen metabolite repression-insensitive manner ^38^. In contrast, no evidence was found for an enzymatic route for NO production in *Magnaporthe oryzae* during hyphal growth and formation of the pathogenic structure appressorium, implying the existence of an alternative NO synthesis pathways in this fungus ^42^. In this study, NO production was abolished while *NR* expression reduced in FgANK1 and FgZC1 mutants, suggesting that these proteins directly or indirectly affect *NR* expression and NO production, a possibility that requires future investigations.

The observation that both FgANK1 and FgZC1 mutants are deficient in NO production and pathogen virulence, suggests a causative link between these two fungal processes. Indeed, virulence was compromised in the rice blast fungus *M. oryzae* when rice leaves were treated with a NO inhibitor at the time of inoculations ^42^. However, possible effects of NO inhibitors on plant defense are unknown and in the absence of a NO-biosynthesis mutant, the link between NO and fungal virulence remains somewhat tenuous.

In addition to the enabling of the discovery of NO being an essential fungal signal following host-sensing, our transcriptome analyses showing distinct gene expression profiles from host-recognition to fungal colonization revealed new insights into the processes involved in root infection by *Fg.* Our results indicated that in response to root exudate-derived signals, the fungus launches an initial ‘preparation-for-infection’ stage presumably via NO signaling. During this stage, several virulence factors were down-regulated while fungal defense and cellular reorganization processes were induced. Suppression of genes involved in DON biosynthesis is particularly interesting given that DON is a virulence factor during wheat head infection in *Fg* ^59^ and root-infection in the related species *Fusarium pseudograminearum* ^60^. *Tri5* expression was also suppressed by SNAP (Supplementary Fig. 3 and 6b), suggesting that host-sensing mediated NO production is responsible for the suppression of toxin biosynthesis genes. It can be speculated that *Fg* might benefit from transcriptional suppression of these virulence factors during the pre-contact stage to avoid host recognition. Pre-contact responsive genes we identified may be closely associated with sensing of environmental signals and host recognition and preparation for infection. Indeed, independent knock out mutants we generated for five cell surface or membrane-associated genes (*FG05_02447*, *FG05_02961*, *FG05_11577*, *FG05_11507* and *FG05_06479*) likely associated with sensing environmental cues showed reduced virulence in *Bd* roots (Supplementary Fig. 4 and Table 2).

Simultaneous upregulations of *Fg* genes encoding transporters of both preferential and non-preferential nitrogen sources at the pre-contact stage were particularly intriguing (Supplemental data 2, 7 and 10). In *Fg* and other fungal species, N transporters and permeases, along with many metabolic enzymes, are tightly regulated by orthologs of the N catabolite regulators AreA/Nit2 ^61–65^. Accordingly, strong expression of non-preferential N assimilation genes such as nitrate and urea transporters is observed under N limitation ^61,64^. As mentioned above (Supplemental data 4), we did not observe a changed expression of the *Fg* AreA homolog in our pre-contact DE gene set. We note that TFs such as AreA are normally upregulated during N starvation. The pre-contact media we used contains ammonium nitrate and should be rich in nutrients (e.g. N compounds) because of the metabolites/exudates produced by the roots. Therefore, the function of AreAs may not be required under the experimental system used here.

In conclusion, the results presented here are consistent with a model (Fig. 8) where sensing of host-derived signals prior to host contact alters diverse processes and prepares the fungus for colonization. NO production, an early response to host-sensing, is regulated by FgANK1 in collaboration with FgZC1 in *Fg*. NO, in turn, can act as a versatile signal potentially involved in fungal development and virulence-associated processes by transcriptionally regulating downstream responses. Future research is required to determine if similar processes are operational in other soil-borne pathogenic fungi but the presence of FgANK1 and FgZC1 homologs in a number of pathogenic fungi and especially in *Fusarium* spp. suggests that this process might be conserved. The new insights revealed by this work will be useful for the development of new disease control strategies that can directly or indirectly target NO production in pathogenic fungi.

## Methods

### Bacteria, fungal and plant material and growth conditions

The *Escherichia coli* strain Top10 (Life Technologies) and *Saccharomyces cerevisiae* strain BY4743 were used for cloning purposes. The *S. cerevisiae* strains Y187 and Y2HGold (Clontech) were used for yeast two hybrid assays. The *Fg* CS3005 isolate was used in growth and infection assays and for generating gene deletion mutants. *Fg* strains were either routinely maintained on Potato Dextrose Agar (PDA, BD Difco) or grown on minimum medium (MM) ^66^ at pH 7 adjusted with MES. For fungal stress test, growth was assayed on PDA plates supplemented with 0.7 M NaCl, 2.2 mM H_2_O_2_ and 0.5 mg/mL calcofluor white. *Bd* (Bd21-3) seeds were surface sterilized and pre-germinated for 5 days prior to *Fg*-roots interaction assays conducted on MM.

### RNAseq and transcriptomic analyses

Fungal and root materials were frozen in liquid nitrogen and RNA was extracted using a QIAgen (Melbourne, Australia) RNeasy spin plant RNA extraction kit according to the manufacturer’s instructions. Four technical replicates consisted of mock or pre-contact fungal mycelium excised from separate plates, or 10-12 infected plants, were pooled. RNA sequencing was performed using three biological replicates with 50 bp single-end on an Illumina HiSeq2500 High Output platform by the Australian Genome Research Facility (Melbourne, Australia). Transcript abundance and differential gene expression were quantified according to the method described previously ^67^. Briefly, quality of sequence reads was assessed by FastQC prior to alignment. By using Tophat2 and Bowtie2 with default settings, all reads were first filtered against the annotated *Bd* genome (v3.1) and all unmapped reads were then mapped to the annotated *Fg* genome ^68^. Subsequently, the transcript abundance was geometrically normalized by Cuffdiff and measured as FPKM (fragments per kilobase of gene model per million reads mapped) values. To assess variation of the samples, unsupervised principal analysis was performed for mean normalized FPKM values using the R packages CummeRbund and ggplot2. Individual sample files were merged for pairwise comparison (fungal alone vs pre-contact vs colonization) using Cuffmerge and analysis of differential expression of genes was performed using CuffDiff. For expression analysis, exon read counts with multiple alignment were not considered. Genes with a |log2 fold change| ≥ 1 and Benjamini and Hochberg-adjusted P value < 0.05 were considered differentially expressed.

### Annotation and functional categorization of differentially expressed genes (DEGs)

Peptide sequences encoded by all DEGs across pairwise comparisons were aligned by BLASTP against a local protein database (-max_target_seqs 1, -evalue 1×10^-5^) generated using the UniProt uniref90 (201706 released) and tabular output was loaded into Blast2GO graphical interface. BLASTP reciprocal best hit analyses were performed in order to identify putative orthologous genes and match unique gene identifiers of the *F. graminearum* CS3005 and PH-1 strains ^68^. To classify stage-dependent transcriptional profiles, fuzzy clustering was performed for the DEGs using the Mfuzzy Bioconductor package ^69^ in R (www.r-project.org). Number of clusters was selected based on the possibility of expression pattern across the tested growth stages of the fungus. GO enrichment analyses for gene clusters were performed with the TopGO Bioconductor package in R using weight algorithm with Fisher’s exact test and an overrepresented P value cutoff of < 0.05 ^70^. Based on the PH-1 identifiers, DEGs from all pairwise comparisons were functionally categorized according to the Database for Annotation, Visualization and Integrated Discovery (DAVID, v6.8) functional catalogue ^71^. Functional categories were considered as enriched in the genome if an enrichment Benjamini-Hochberg adjusted P value is smaller than 0.05.

### Annotation of specific gene categories

Protein cellular localizations were predicted using the BUSCA web server with multi-software integration ^72^. Secretome of *Fg* was inferred from a previous report ^73^. Putative effector proteins were predicted using EffectorP2 with default settings ^74^. To identify secreted peptidases, sequences of the predicted secreted proteins were subjected to a local BLASTP database based on MEROPS ^75^. Transporters were identified and classified by BLAST searching against the Transporter Collection Database (http://www.tcdb.org/). To predict carbohydrate-active enzymes, sequences of DEGs were aligned against the Carbohydrate-Active enZYmes Database (<u>CAzyme</u>) using HMMER3 (e-value < 1×10^-17^ and coverage > 0.45). Enzyme capacity was allocated to the predicted CAzymes as described before ^76^. For lipase prediction, HMMER3 search (e-value < 1×10^-17^ and coverage > 0.45) for the DEGs against the Lipase Engineering Database was performed ^77^. Genes encoding key secondary metabolism enzymes, transcription factors, kinases and G-protein coupled receptors were matched to the previous annotation in the PH-1 strain ^22,78–80^.

### Quantitative real-time PCR analysis

Total RNA was treated with DNase (Qiagen) and 0.5-1 µg was prepared for first-strand cDNA synthesis using the superscript synthesis kit (Invitrogen, USA) and quantitative real-time RT-PCR (qRT-PCR) was performed using the ViiA 7 real-time PCR detection platform (Applied BioSystems). All gene specific primers used for qRT-PCR were evaluated by BLASTN search against the *Fg* and *Bd* genomes to exclude unspecific annealing and are listed in Supplemental Table 2. Expression levels were normalized to the fungal house-keeping gene *α*-*Tubulin* and were averaged over at least three biological replicates.

### Vector construction and fungal genetic manipulation

Targeted deletion in *Fg* was based on homologous recombination described previously ^81^. Genomic region of each gene was replaced with corresponding deletion cassettes and subsequent mutants were identified by PCR assays with relevant primers (Supplemental Table 2). Deletion vectors were constructed using pYES2 plasmid as a backbone by homology dependent assembly in yeast ^82^. For these assemblies, double-stranded DNA synthesis (gBlocks, IDT) or PCR amplifications were implemented to generate the following fragments: 940 bp sequence region immediately upstream of the target genes, a *gpdA* promoter-*npt* cassette conferring fungal geneticin resistance ^81^, and 940 bp sequence region immediately after the stop codon of each gene. The upstream and downstream sequences both harbor additional 30 bp homologous overhangs at their 5 and 3 prime ends (sequences as indicated in the primers in Supplemental Table 2). The above fragments along with the *Hind*III/*Xba*I linearized backbone were introduced into yeast ^83^. Clones were verified by yeast colony PCR using specific primer pairs (Supplemental Table 2), recovered using a Zymoprep Miniprep II kit (Zymo Research, USA) and subsequently transformed into *E. coli* for amplification. To generate the *GFP*-*FgANK1* fusion used for complementation and protein localization assays, the following fragments were synthesized and assembled in yeast: a 1870 bp above mentioned *gpdA* promoter fragment or 970 bp sequence region immediately upstream of the *FgANK1* start codon each with an additional 30 bp 5’ end homologous tag sequence, a 1816 bp *GFP* (without stop codon)-10×Glycine linker-*FgANK1*-CDS cassette containing a 30 bp 5’ end overhang homologous to the upstream promoter sequences, and a 1706 bp *TrpC* promoter-*nat*-*TrpC* terminator cassette concatenated by a 940 bp sequence behind the stop codon of *FgANK1* with a 30 bp sequence tagging the vector backbone. The *gpdA* and native promoter cassettes were introduced into the ΔFgANK1 strain, and mutants were selected with 50 mg/L nourseothricin. Using the same strategy, *TagBFP* fusion cassettes were constructed with a 699 bp *TagBFP* fragment without stop codon. For the ANK repeats and VWA domain truncations, sequences coding the FgANK1^16–235^ and FgANK1^247–484^ regions were deleted, respectively. To generate *FgZC1*-*HA*-*nptII* cassette fusion, a 970 bp genomic region including the upstream and the first 30 bp region of *FgZC1* and a 2834 bp sequence consisting of the *FgZC1* cDNA (without stop codon) linked with a 54 bp 8×Glycine linker-*HA*-stop codon sequence followed by a 668 bp *gpdA* terminator and a 30 bp 3’ end overlapping with the *nptII* cassette were synthesized and used for assembly. For co-immunoprecipitation assays, the FgZC1-HA strain was crossed with the Pro_FgANK1_–tagBFP-FgANK1 and Pro_FgANK1_–tagBFP-FgANK1^ΔVWA^ strains as described before ^84^. The resulted recombinant progeny could grow on PDA with both 50 mg/L nourseothricin and 50 mg/L geneticin and be immunoprecipitated by Western blot. Vectors used for yeast two-hybrid (Y2H) analyses were constructed and introduced into the yeast strain Y2HGold according to manufacturer’s protocol (Clontech). The *FgANK1* cDNA regions were amplified using the primers 5Arm_02877F, 3Arm_02877R, Y2HANK-R and Y2HVWA-F (Supplemental Table 2) and corresponding deletion vectors as templates confirmed by sequencing.

### Yeast two-hybrid library construction, screening and protein-protein interaction analyses

Y2H library was constructed using the Make your own “Mate & Plate” library system according to manufacturer’s manual with minor modifications (Clontech). In brief, fungal mycelia during the pre-contact stage were flash-frozen and processed for RNA preparation. Two μg of DNase-treated total RNA was used for first strand cDNA synthesis. The resulted cDNA was PCR enriched using Phusion polymerase (NEB) with cycling parameters: 98 °C for 30 s followed by 24 cycles of 98 °C for 10 s, 68 °C for 3 min (extension time was increased by 3 s with each successive cycle) and 68 °C for 5 min. Subsequently, the PCR products were size-selected and purified to generate a pool of high quality double stranded cDNA, of which a total of 10 μg was co-transformed with *Sma*I linearized pGADT7^rec^ into the haploid strain Y187. Transformation mixture was spread out on 100 synthetic dropout (SD)/-Leu agar plates (150 mm diameter, Corning). Transformants were then pooled and prepared as 1 ml aliquots. For library screening, two independent experiments were performed by mating the aliquoted cells with the Y2HGold strain harboring the bait construct pGBKT7-*FgANK1* following instruction of the Matchmaker Gold yeast two-hybrid system (Clontech). Interaction clones were selected on 50 SD/-Trp/-Leu/X-α-Gal/AbA (40 μg/ml X-α-Gal and 200 ng/ml Aureobasidin A) agar plates. Further selection was conducted for the candidate colonies by segregating them on high-stringency plates SD/-Ade/-His/-Trp/-Leu/X-α-Gal/AbA for two generations. To exclude false positives, plasmids were recovered from the screened candidate preys and co-transformed with the bait vectors into Y2HGold cells plating under both low- and high-stringency. The confirmed clones were considered as positive interaction partners and sequenced. Protein-protein interactions were assayed by co-transforming the bait vectors pGBKT7-*FgANK1*, pGBKT7-*FgANK1*^Δ^*^ANK^*, and pGBKT7-*FgANK1*^Δ^*^VWA^* with the prey vector pGADT7-*FgZC1* in pairs. Control vectors provided in the kit (Clontech) were included to compare positive and negative interactions and to test self-activation of both baits and prey.

### Protein extraction, Western blot and co-immunoprecipitation (co-IP) assays

For protein extractions, fungal strains were conditionally grown on polyvinylidene fluoride (PVDF) membranes (GE Healthcare) placed on MM plates. To prepare cytosolic fractions, fungal mycelia collected from ten plates were suspended in 20 ml 1M sorbitol solutions containing 500 mg driselase and 200 mg lysing enzymes (Sigma). The solutions were incubated at 28 °C, 70 rpm for 30 min and centrifuged at 1800×g, 4 °C. Cell pellets containing protoplasts were resuspended in 5 mL lysis buffer (250 mM sucrose, 20 mM Tris-HCl [pH 7.4], 1 mM EDTA, 5 mM DTT, and 50 µL protease inhibitor cocktail [Sigma]) and homogenized on ice. The homogenates were filtered through 50 µm nylon meshes (Biologix) and spun at 1800×g for 10 min. The supernatants were subsequently centrifuged at 10,000×g for 10 min at 4°C and collected as cytosolic fractions. The pellets were resuspended in sucrose cushions and subjected to nuclear fractionation as previously described ^85^. Cytoplasmic and nuclear protein extracts were separated using pre-casted Bolt 4–12% Bis-Tris gels and transferred to PVDF membranes followed by protein detection, all according to manufacturer’s instructions (Invitrogen). Proteins were detected using polyclonal antibodies against GFP (Roche), Histone H3 (GeneTex) (nuclear control), and a monoclonal antibody against GAPDH (GeneTex) (cytoplasmic control). For total protein extraction, fungal mycelia were thoroughly ground in liquid nitrogen and suspended in 5 volumes PBS buffer containing 1% Triton X-100 and 1/10 volume protease inhibitors cocktail. The extracts were centrifuged at 14000×g for 20 min at 4 °C and the supernatants were collected as total protein extracts. Co-IP assays were performed by incubating total proteins with anti-HA (Sigma) agarose followed by column wash and elution according to manufacturer’s instructions (Pierce, Thermo Fisher Scientific). The protein samples were detected with a polyclonal anti-tRFP antibody (Evrogen) for TagBFP, and the monoclonal anti-GAPDH and -HA antibodies as loading controls

### Plant infection assays

Fusarium head blight (FHB) assays were conducted on wheat cultivar cv. Kennedy. Ten µL of 10^6^ spores/ml *Fg* macroconidia harvested from liquid carboxymethylcellulose (CMC) medium were drop-inoculated at the fifth spikelet from the base of the inflorescence of each spike. The infection was maintained at 25°C with fluorescent lighting provided for 16 hours per day with 60% humidity. Inoculated heads were immediately covered with a humidified plastic zip lock bag which was then changed by a glassine bag at 3 days post inoculation (dpi). Heads with typical symptoms were examined at 10 dpi and scored by counting the total number of spikelets showing symptoms. For *Fg*-*Bd* root interaction assays, fungal strains were transferred from CMC agar to center or bottom of MM plates and pre-grown for 3 days to allow approximately 2 cm of proliferating hyphae. Five-day-old *Bd* seedlings were then distantly placed above the fungal colonies. Fungal mycelium derived from the center-placed plugs reached to *Bd* roots at 3 dpi and caused visible root necrosis at 5 dpi (colonization). At this point (5 dpi), the fungal mycelium from the bottom of the plates were still growing towards the roots (pre-contact). The roots and fungal mycelium were covered by aluminum foil to maintain dark and plates were placed in a slanting position (with an angle of approximately 45°) in a growth chamber (Conviron) using a 16 /8 h light/dark cycle. Samples were collected at 5 dpi for RNA extraction. Root disease symptoms were recorded for the infected roots at 7 dpi and indexed as percentage of root necrosis. Inoculated or mock plates were slightly opened to prevent volatile accumulation. In most cases, two independent mutants for each gene deletion with two replications were used.

### Root and fungal exudate extraction

To prepare root exudates, approximately 15 mL of plant-alone and pre-contact medium from 5-day-old plates were flash frozen and freeze-dried overnight. Five plates of each treatments were used. Samples were suspended in 5 mL of extraction solution (Ethyl acetate: methanol: dichloromethane: water 3:2:1:2, v/v) and sonicated for 15 min followed by brief centrifugation. After filtration through 0.45 µm filters (Millipore), the supernatants were dried under nitrogen and resuspended in methanol: water (1:1, v/v). Each treatment was pooled to generate a final volume of 200 µL and used as metabolite extracts to treat the fungus in vitro.

### Staining, fluorescence quantification and microscopic analyses

To observe nitric oxide (NO) production, fungal hyphae were stained using 10 µM fluorescent dye 4-amino-5-methylamino-2,7-difluorofluorescein diacetate (DAF-FMDA) (Cayman Chemical). To confirm genuine reaction between DAF-FMDA and NO, a cell-permeant NO scavenger 2-(4-carboxyphenyl)-4,4,5,5-tetramethylimidazoline-1-oxyl-3-oxide (cPTIO) (Sigma) at a high concentration of 500 μM was included. DAF-FMDA combined with or without cPTIO was directly applied to the edge of the mycelium towards roots or the same location on mock plates. For kinetic assays, 96-well plates with 10^4^ conidia/well in 100 µL MM supplemented with combinations of pre-contact metabolite extracts, DAF-FMDA and cPTIO were used. Signals were detected with GFP filter sets in a 48-hr fluorescence kinetics by a BioTek Cytation1 imaging reader and the florescence was subtracted and corrected from background. Three biological replicates were measured per sample and the experiments were independently repeated twice. Combinations of FITC, DAPI and Rhode filter sets equipped on a Zeiss Axio Imager M2 microscopy were used to observe GFP and BFP protein markers as well as DAF-FMDA, DAPI (Sigma) and FM 4-64 (Invitrogen) staining, and a Carl-Zeiss ZEN software (v. 2.3) was used to record florescence signals and intensities.

## Data availability

RNA-seq data have been deposited in the NCBI BioProject database with accession code PRJNA564465. Data underlying Figs. 2, 3h, 4e, 5d, 6b, 7f and Supplementary Figs. 1, 2, 3, 4, 5d, 6a, 6b, 8g, 9b and 9c are provided as Source Data files in Supplementary Data 11. All other data for this paper are available from the authors upon reasonable request.

## Acknowledgements

YD was the recipient of a CSIRO Research Office post-doctoral fellowship. We thank Drs Melania Figueroa and Jonathan Anderson (both from CSIRO Agriculture and Food) for their suggestions of the manuscript.

## Contributions

Y.D, D.M.G and K.K designed the experiments. Y.D. performed the experiments and analyzed the results. Y.D and D.X analyzed the RNA-seq data. Y.D, D.M.G and K.K wrote the manuscript. All authors read and approved the final manuscript.

---

**Figure S1.** The top-two induced putative effector genes *FG05_00009* and *FG05_11513* at the pre-contact stage may regulate stress responses and virulence in *Fg.* **a** Growth phenotypes of *Fg* strains were tested on PDA plates supplemented with 2 mM H_2_O_2_ (oxidative stress), 1 M NaCl (osmotic stress), or 0.7 mM Calcofluor (cell wall stress). **b**-**c** Pathogenicity assays of the mutant *Fg* strains towards wheat heads and *Bd* roots. Photographs show representative infection data from one of the two independent mutants generated for each gene.

**Figure S2.** A role for putative urea (DUR) transporters of *Fg* in fungal development and virulence. **a** Nitrogen assimilation assays for WT and DUR mutants on MM supplemented with glucose and different nitrogen sources. **b** Growth phenotypes of WT and DUR3 and DUR31 mutants were tested on PDA plates supplemented with 2 mM H_2_O_2_ (oxidative stress), 1 M NaCl (osmotic stress), or 0.7 mM Calcofluor (cell wall stress). **c** Pathogenicity assays of DUR3 and DUR31 mutants on wheat heads and *Bd* roots. Photographs (left panels) show representative infection data from the two independent mutants generated for DUR3 and DUR31, respectively. Percentage *Bd* root necrosis and percentage spikelets infected (right panels) are averages from a minimum of twenty replicate roots and at least five wheat heads, respectively, with error bars representing the standard deviation (One-way ANOVA with Dunnett’s multiple comparisons test, **P< 0.01, ***P< 0.005, ****P< 0.001).

**Figure S3.** NO differentially regulates the expression of different pre-contact genes in *Fg.* Expressions of six pre-contact DEGs in response to an external NO donor (SNAP) or NO scavenger (cPTIO) were analyzed in *Fg* grown in liquid MM supplemented with 2% glucose for three days. 300 µM SNAP and 500 µM cPTIO were then individually or together supplemented to the pre-grown cultures. 0.001% ethanol solvent was used as mock treatment. After 6 h, the fungal mycelia were harvested and frozen in liquid nitrogen and immediately subjected to RNA preparation and first strand cDNA synthesis for RT-qPCR experiments. Gene expression levels were normalized using the fungal reference gene α*-Tubulin*. Data are the mean of three biological replicates with error bars representing standard error of the mean. Asterisks represent differences that were statistically significant compared with the mock control (One-way ANOVA with Fisher’s Least Significant Difference test, *P< 0.05. **P< 0.01, ***P< 0.005).

**Figure S4.** PCR confirmation for all *Fg* transformants and pathogenicity phenotypes of the selected pre-contact DEG deletion mutants. For the positive targeted deletion mutants, multiplex PCR amplification only generated a single band with a size that differs from that of the WT. Amplification of the ectopic strains resulted in two bands representing WT loci and the deletion constructs. Growth phenotypes on PDA agar (left to right: no supplement, supplemented with 2 mM H_2_O_2_, 1 M NaCl, and 0.7 mM Calcofluor) were recorded at 7 days and the representatives show similar results from two independent mutants with two replicates. Fungal pathogenicity on *Bd* roots (7 dpi) and wheat heads (10 dpi) was compared to the CS3005 WT strain. Where significantly reduced virulence was observed (blue bars), two independent deletion mutants were shown (One-way ANOVA with Dunnett’s multiple comparisons test, *P< 0.05, **P< 0.01, ***P< 0.005, ****P< 0.001). Mutant virulence in wheat head was not considered to be affected if any individual measurement is above the base line.

**Figure S5.** FgANK1 is required for host-sensing mediated NO production but not for fungal growth on PDA in *Fg*. **a** Growth rates of WT, ΔFgANK1, and complemented strains of ΔFgANK1 on PDA agar plates. No significant difference in growth could be observed among the tested strains. **b** FgANK1 is required for pathogenicity on *Bd* roots. No symptom development was evident when *Bd* roots were directly inoculated with the ΔFgANK1 mutant for three weeks post-inoculation (wpi) while *Bd* plants similarly inoculated with the WT *Fg* were nearly dead after two weeks. **c** FgANK1 is required for host-sensing mediated NO production in *Fg*. ΔFgANK1 showed abolished NO production at the pre-contact stage (upper panel) or after treatment with pre-contact root metabolites (lower panel). **d** Expression of FgANK1 was not regulated by external NO or NO scavenger cPTIO in both WT *Fg* and FgANK1 complemented strains.

**Figure S6.** FgANK1 likely acts upstream of NO production in *Fg*. **a** Fungal marker genes were not responsive to pre-contact metabolites but responsive to external NO donors in the ΔFgANK1 mutant. Before the treatment, the mutant was grown in liquid MM supplemented with 2% glucose for three days. Expressions of fungal genes differentially regulated by root pre-contact under conditions when the ΔFgANK1 mutant was treated with media extracts from plant alone, fungal-plant pre-contact and solvent solution (methanol and water 1:1). No statistically significant differences were found for transcript levels of the selected genes among different treatments. **b** Expressions of fungal genes differentially regulated by root pre-contact when ΔFgANK1 was treated for 6 hrs with 300 µM SNAP, 500 µM cPTIO or both. 0.001% ethanol solvent was used as mock treatment. No difference in gene expressions was found for any of the tested genes in comparison between the ΔFgANK1 mutant and its complemented strain Pro_FgANK1_::GFP-FgANK1. Except for *FG05_01947*, which encodes a nitrate reductase (NR), the other five tested genes showed similar regulation patterns to the pre-contact stage regardless of FgANK1. Thus, FgANK1 acts upstream of NO production in *Fg*. Gene expression levels were normalized using the fungal reference gene *α-Tubulin*. Data are the mean of three to four biological replicates with error bars representing standard error of the mean. Asterisks represent differences that were statistically significant compared with the mock control (unpaired two-tailed *t*-test, *P< 0.05, **P< 0.01, ***P< 0.005)

**Figure S7.** Homologs of FgANK1 in different fungal and eukaryotic species. **a** FgANK1 (489 aa) was aligned against other sequences found in the STRING database and the top hits were displayed. Grey lines represent similar protein sequences anywhere in STRING. Sequence similarity was interpreted as alignment bitscore of similar proteins in a taxon of choice. **b-c** Phylogenetic analysis of ankyrin-domain containing proteins (**b**) and FgZC1 homologs. The analyses were conducted by customized ete3 reconstruction pipeline ^86^. The protein evolutionary models were tested using Neighbor-Joining inference and then referred to the tree building based on Maximum Likelihood method. Amino acid sequences from selected species were obtained from the Joint Genome Institute (JGI) genome portal (http://genome.jgi.doe.gov/). The Arabidopsis proteins ANK1 (AT5G02620) and AtTN1 (AT3G12360) are used and fallen into an outer group. The yeast ASG1 protein sequence was retrieved from the Saccharomyces Genome Database (https://www.yeastgenome.org/). Protein domains found in FgANK1 and FgZC1 are shown schematically in **b** and **c**, respectively.

**Figure S8.** The ankyrin (ANK) repeat domain of FgANK1 is required for virulence towards *Bd* roots but not for fungal growth. Because GFP and NO-detecting DAF-FM DA fluorescence signals were indistinguishable, we constructed TagBFP fusions and introduced into the ΔFgANK1 background. **a** FgANK1 localization is not affected by GFP or TagBFP N-terminal fusions. Microscopic images of 7-day old fungal mycelium and spores expressing Pro*_FgANK1_*::*GFP-FgANK1*, Pro*_gpdA_*::*GFP*-*FgANK1* or Pro*_FgANK1_*::*TagBFP-FgANK1* on the carboxymethylcellulose (CMC) agar medium. Fluorescence signals indicate GFP (left and middle panels) or TagBFP (right panels). **b-f** Compared to the WT (**b**), the FgANK1 mutant strain (**c**) complemented with TagBFP-FgANK1 (**d**), TagBFP-FgANK1^ΔVWA^ (**e**) or TagBFP-FgANK1^ΔANK^ (**f**) showed normal growth under cell wall (Calcofluor), oxidative (H O) osmotic (NaCl) stresses. The expression of the Pro ::*TagBFP-FgANK1*^Δ^*^ANK^* construct in the ΔFgANK1 mutant background could not restore the fungal pathogenicity on *Bd* roots at 7 dpi (**f** right panel and **g**). #2 refers to transformant 2 of two independent transformants generated. Pictures are representatives of two independent fungal transformants (One-way ANOVA with Dunnett’s multiple comparisons test, ****P< 0.001).

**Figure S9.** FgZC1 is required for virulence towards *Bd* roots but not for wheat heads. **a-d** Disease phenotypes and fungal marker gene expression in the ΔFgZC1 mutant. Deletion of *FgZC1* delayed disease development in *Bd* roots (**a** and **b).** *Bd* root necrosis was recorded at 7, 8, 10 and 14 dpi, respectively. The selected fungal pre-contact DEGs showed similar expression patterns in ΔFgZC1 relative to those in ΔFgANK1 when pre-contacted with *Bd* roots (**c**). Transcript levels were normalized using the fungal reference gene α*-Tubulin*. Data are the mean of three to four biological replicates with error bars representing standard error of the mean. Asterisks represent differences that were statistically significant compared with the WT control (One-way ANOVA with Dunnett’s multiple comparisons test, *P< 0.05. **P< 0.01, ***P< 0.005).

**Supplemental Table 1**. Targeted *Fg* deletion mutants characterized in this study.

**Supplemental Table 2**. Primer and gene fragment sequences used in this study.

**Supplemental Data1**. Summary of Illumina sequencing reads mapping to *Fg* and *Bd* genomes.

**Supplemental Data 2**. List of DEGs in the comparison between *Fg* pre-contact and *Fg*-only conditions.

**Supplemental Data 3**. Annotation of all pre-contact DEGs.

**Supplemental Data 4**. List of differentially regulated TF genes during the pre-contact stage of *Fg*.

**Supplemental Data 5**. List of cell wall integrity and stress response component (WSC) domain containing proteins predicted in *Fg*.

**Supplemental Data 6**. Summary of DAVID functional annotation analyses for all *Fg* pre-contact DEGs.

**Supplemental Data 7**. GO enrichment analyses by TopGO for pre-contact DEGs from six clusters.

**Supplemental Data 8**. List of predicted effector genes differentially regulated during the pre-contact stage of *Fg*.

**Supplemental Data 9**. List of SME genes differentially regulated during the pre-contact stage of *Fg*.

**Supplemental Data 10**. List of transporter genes differentially regulated during the pre-contact stage of *Fg*.

**Supplemental Data 11**. Source data.

