## Supplementary Figures 1-9 for "Novel regulators of nitric oxide signaling triggered by host perception in a plant pathogen"

Supplementary Fig. 1

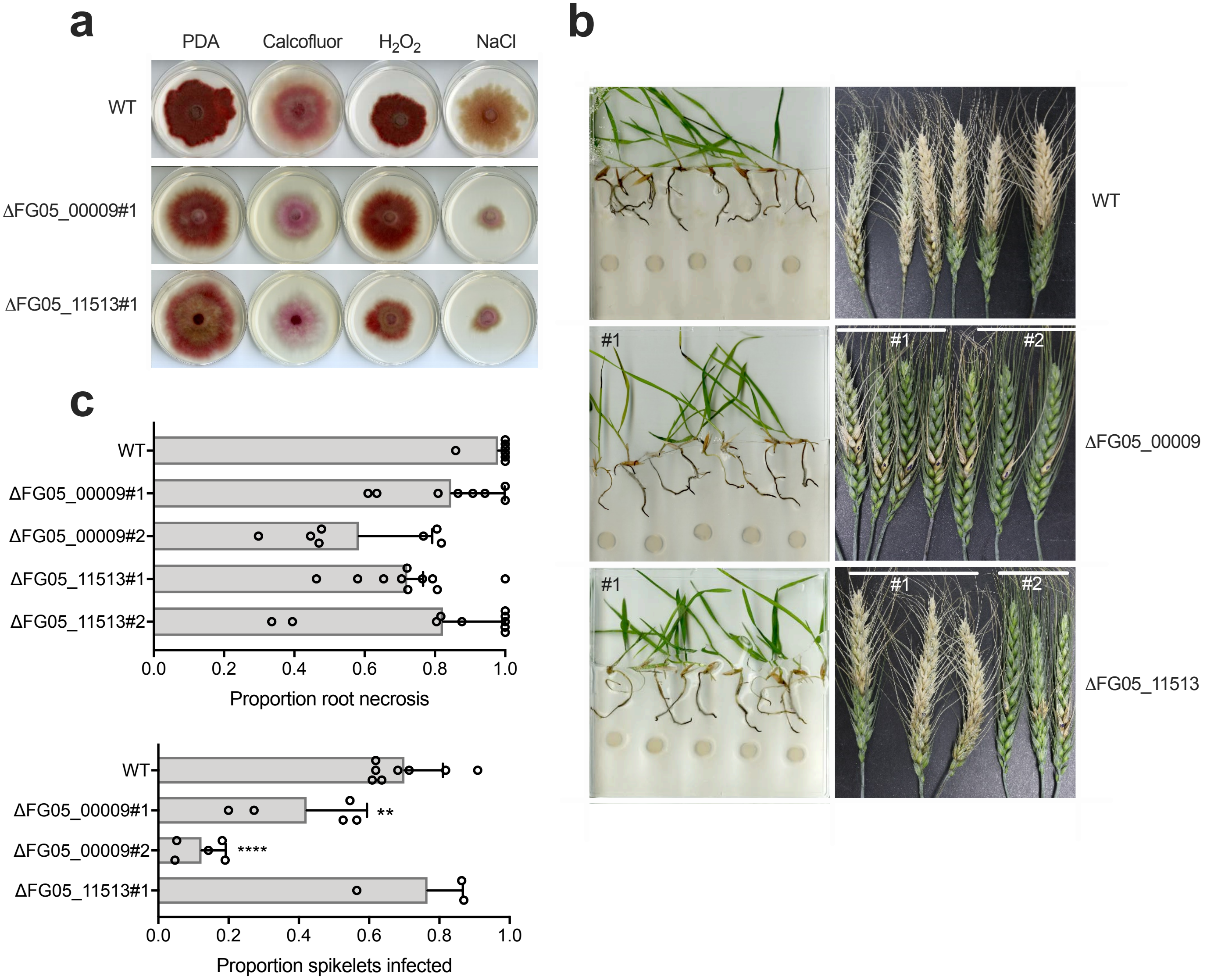

### Suppl. Fig. 2

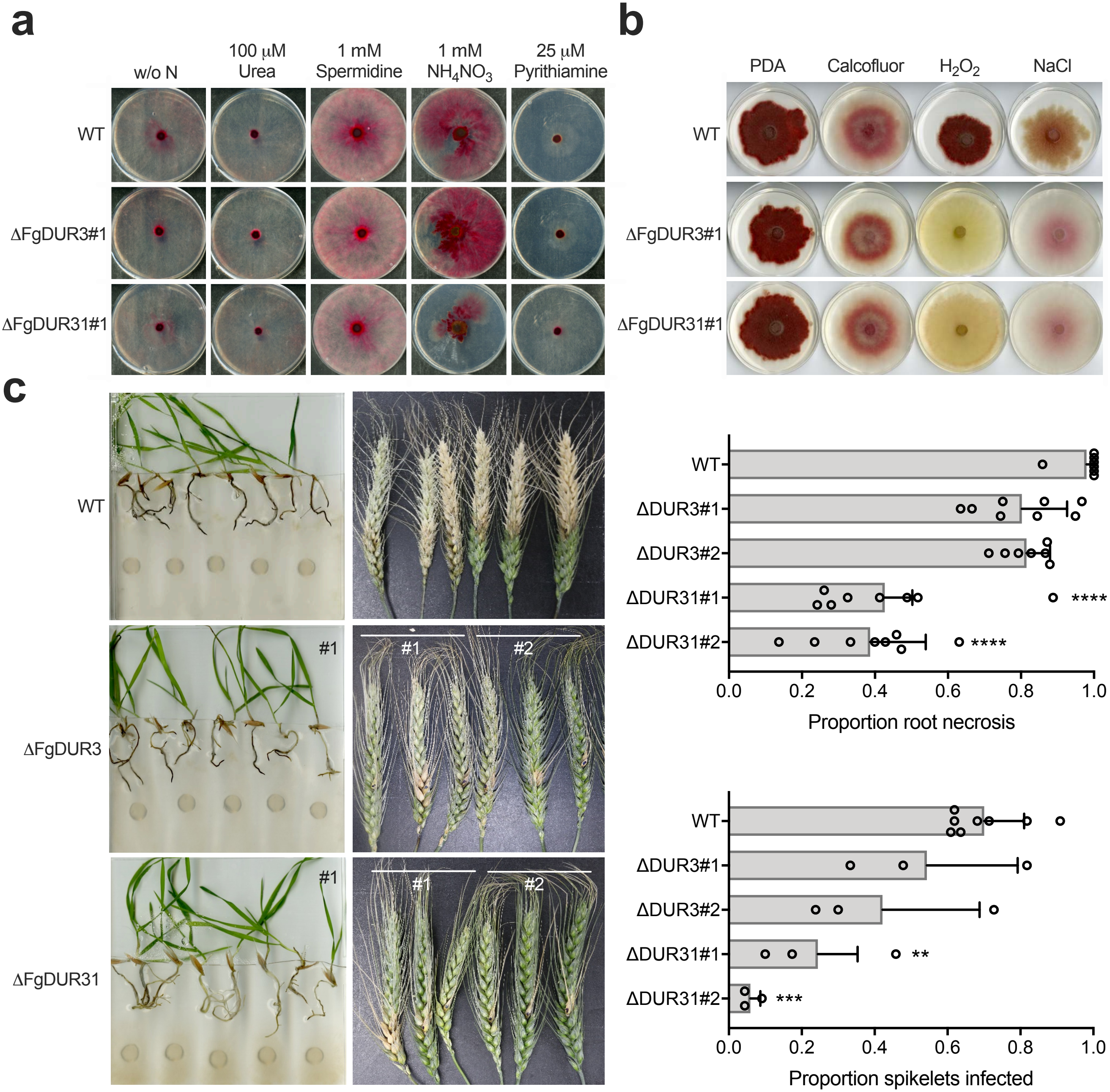

a
w/o N
100 μM Urea
1 mM Spermidine
1 mM $NH_4NO_3$
25 μM Pyrithiamine
WT
ΔFgDUR3#1
ΔFgDUR31#1
b
PDA
Calcofluor
$H_2O_2$
NaCl
WT
ΔFgDUR3#1
ΔFgDUR31#1
c
WT
ΔFgDUR3
ΔFgDUR31
#1
#2
WT
ΔDUR3#1
ΔDUR3#2
ΔDUR31#1
ΔDUR31#2
****
****
0.0
0.2
0.4
0.6
0.8
1.0
Proportion root necrosis
**
***
Proportion spikelets infected

Suppl. Fig. 3

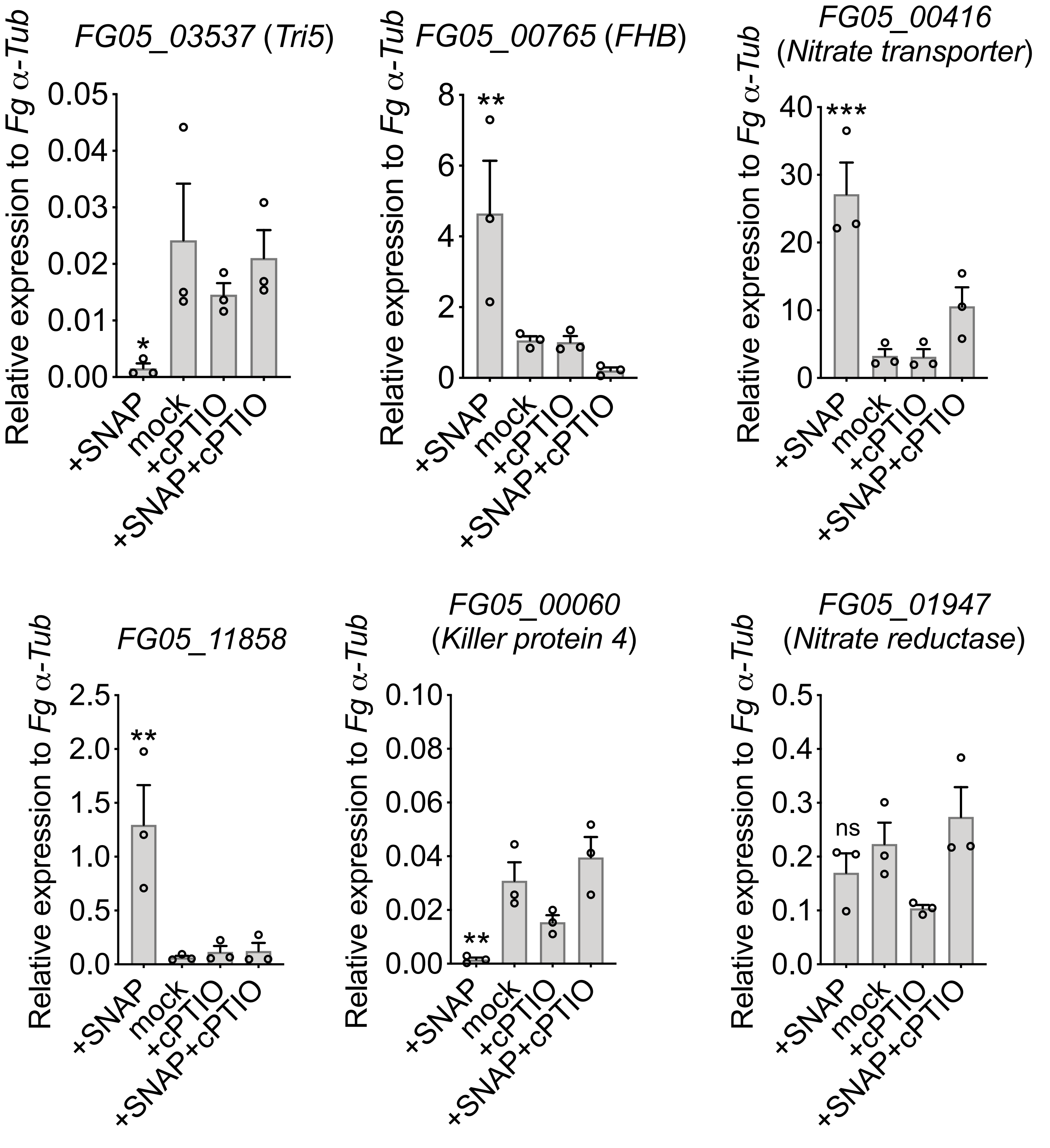

FG05_03537 (Tri5)
FG05_00765 (FHB)
FG05_00416 (Nitrate transporter)
FG05_11858
FG05_00060 (Killer protein 4)
FG05_01947 (Nitrate reductase)
Relative expression to Fg α-Tub
0.05
0.04
0.03
0.02
0.01
0.00
8
6
4
2
0
40
30
20
10
2.5
2.0
1.5
1.0
0.5
0.0
0.10
0.08
0.06
0.04
0.02
0.00
0.5
0.4
0.3
0.2
0.1
*
**
***
ns
+SNAP
mock
+cPTIO
+SNAP+cPTIO

Suppl. Fig. 4

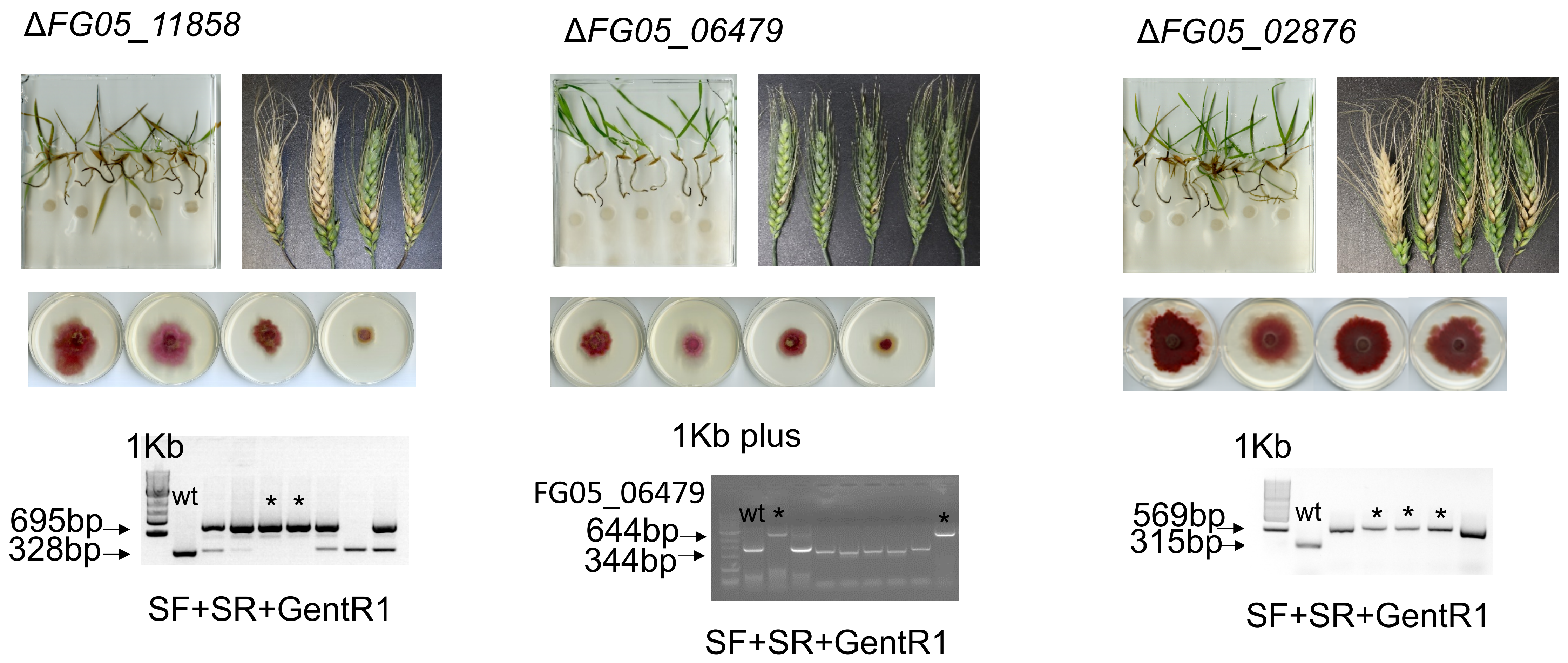
ΔFG05_11858
ΔFG05_06479
ΔFG05_02876
1Kb
wt
* *
695bp
328bp
SF+SR+GentR1
1Kb plus
FG05_06479
wt *
*
644bp
344bp
SF+SR+GentR1
1Kb
wt
* * *
569bp
315bp
SF+SR+GentR1

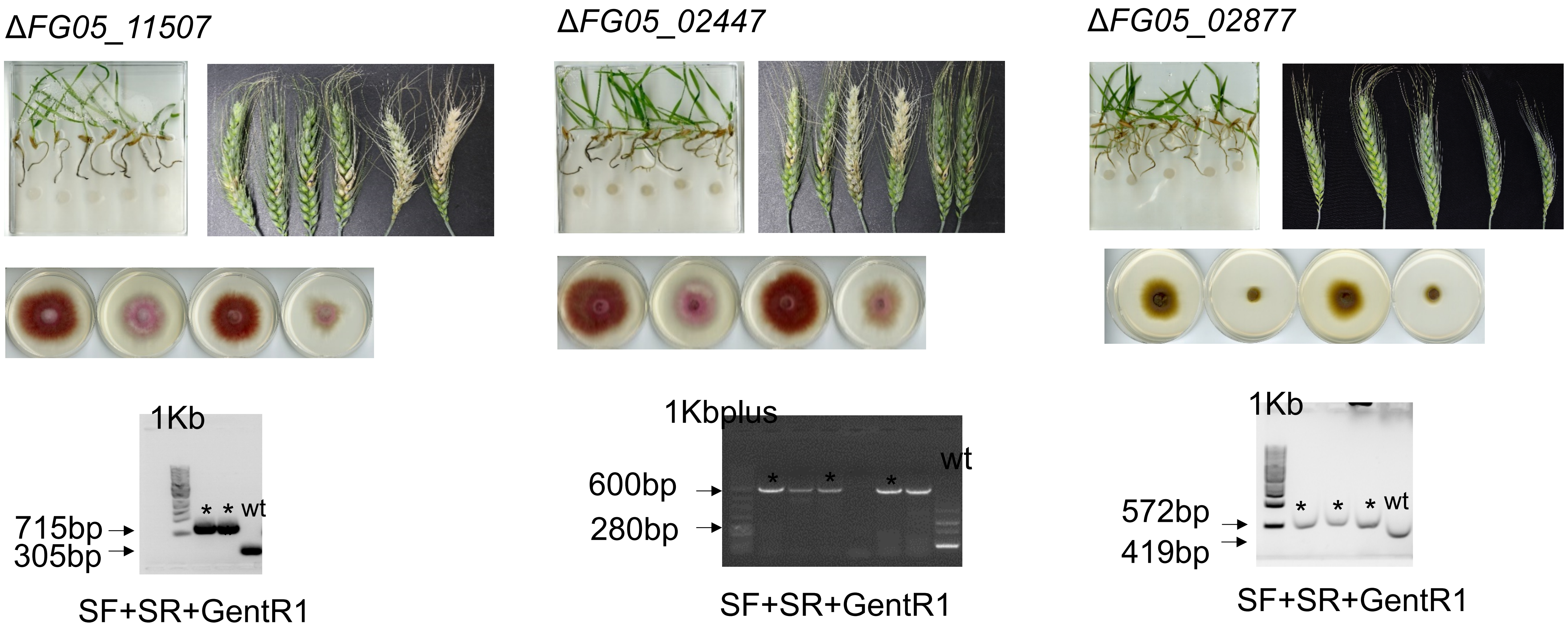
ΔFG05_11507
1Kb
* * wt
715bp
305bp
SF+SR+GentR1
ΔFG05_02447
1Kbplus
wt
600bp
280bp
* * *
SF+SR+GentR1
ΔFG05_02877
1Kb
* * * wt
572bp
419bp
SF+SR+GentR1

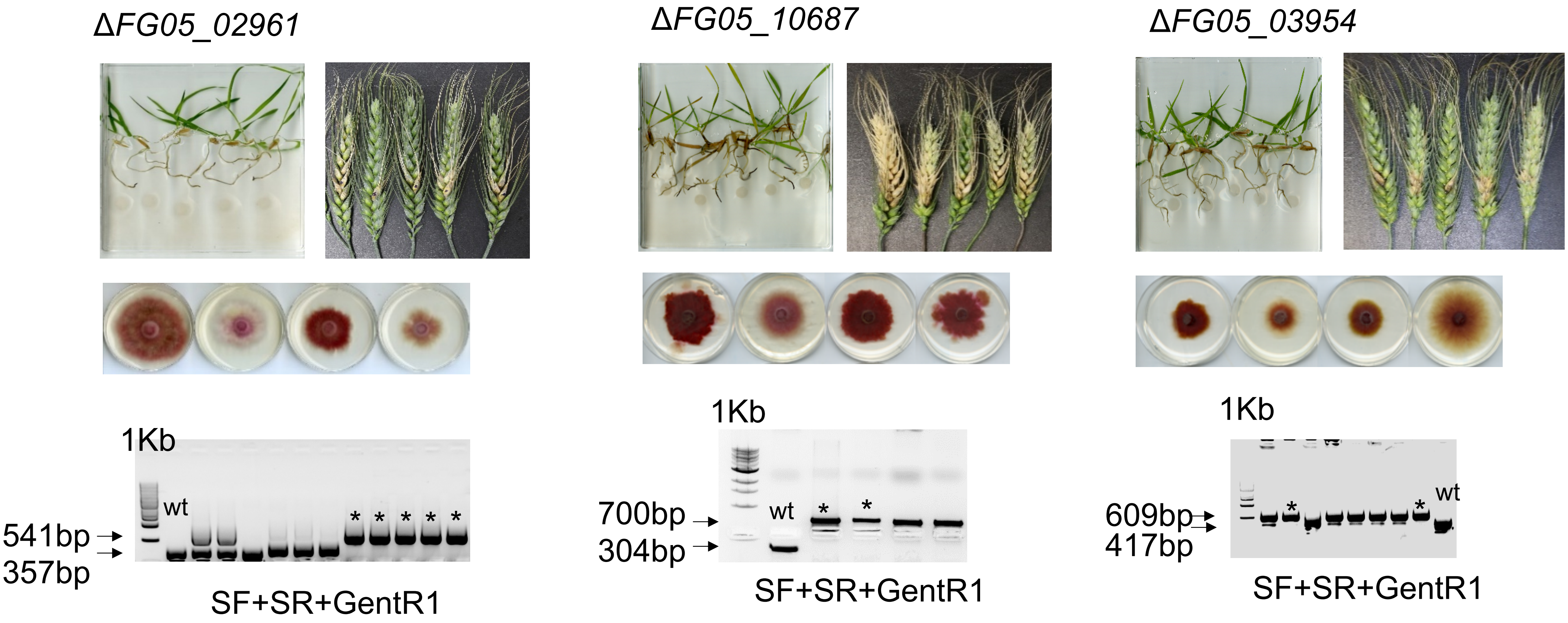
ΔFG05_02961
ΔFG05_10687
ΔFG05_03954
1Kb
wt
541bp
357bp
SF+SR+GentR1
1Kb
wt
700bp
304bp
SF+SR+GentR1
1Kb
wt
609bp
417bp
SF+SR+GentR1

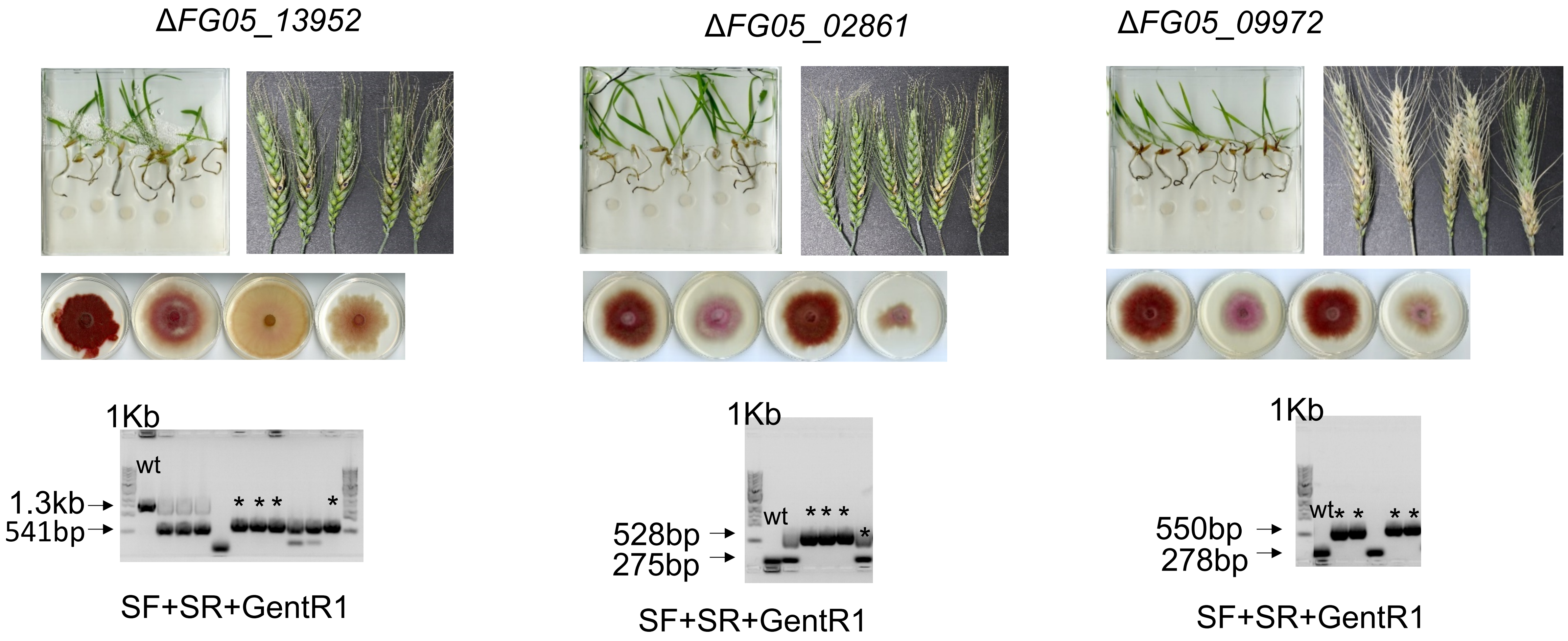
ΔFG05_13952
1Kb
wt
1.3kb
541bp
SF+SR+GentR1
ΔFG05_02861
1Kb
wt
528bp
275bp
SF+SR+GentR1
ΔFG05_09972
1Kb
wt
550bp
278bp
SF+SR+GentR1

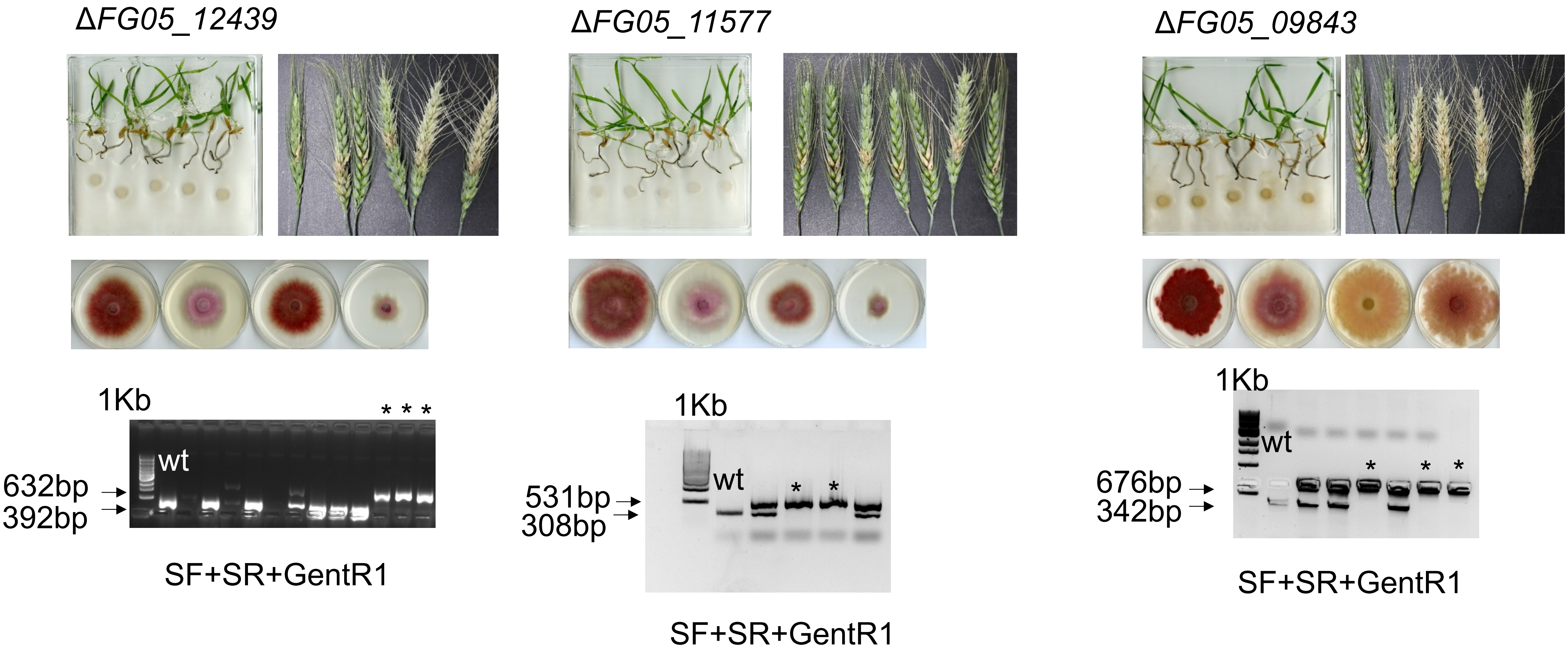

ΔFG05_12439
1Kb
wt
* * *
632bp
392bp
SF+SR+GentR1
ΔFG05_11577
1Kb
wt
* *
531bp
308bp
SF+SR+GentR1
ΔFG05_09843
1Kb
wt
* * *
676bp
342bp
SF+SR+GentR1

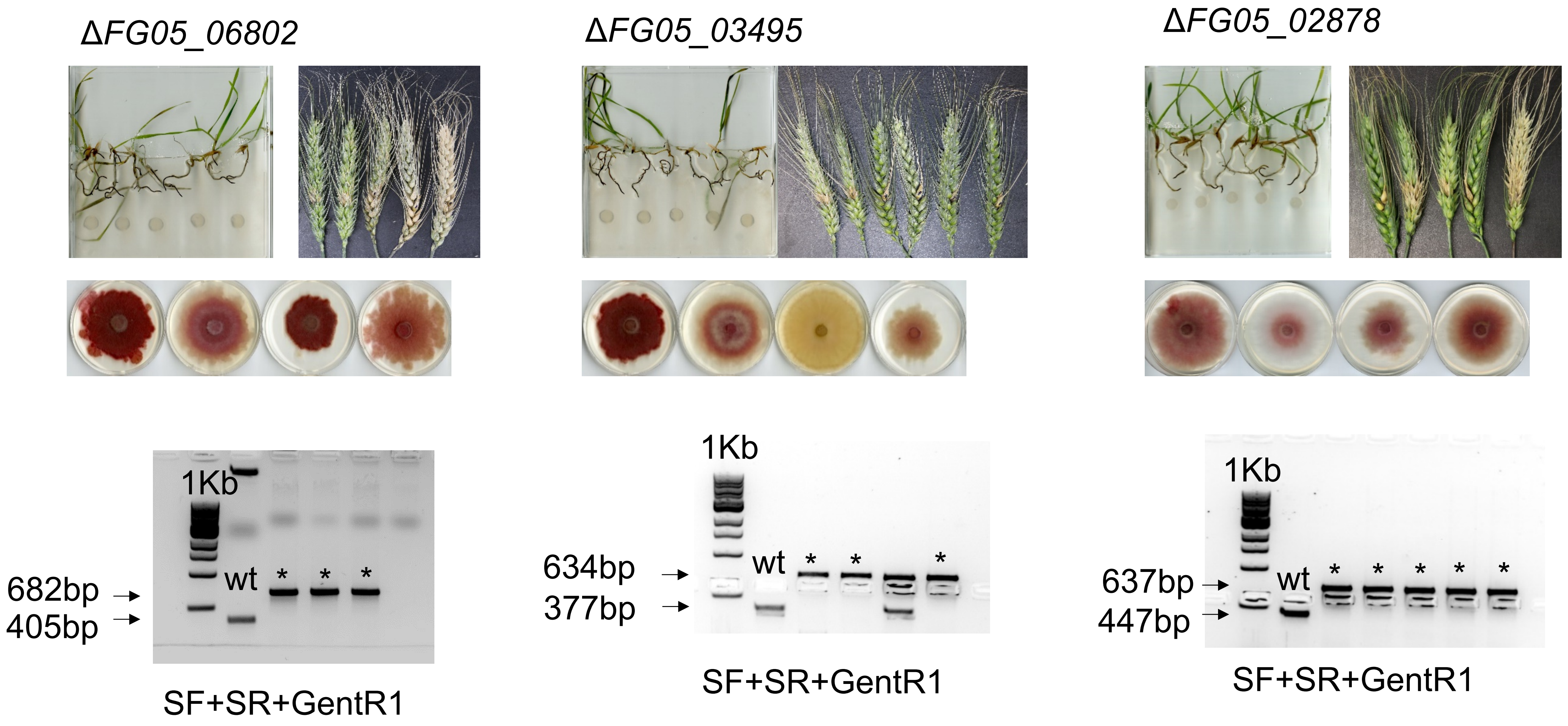

ΔFG05_06802
ΔFG05_03495
ΔFG05_02878
1Kb
wt
682bp
405bp
SF+SR+GentR1
1Kb
wt
634bp
377bp
SF+SR+GentR1
1Kb
wt
637bp
447bp
SF+SR+GentR1

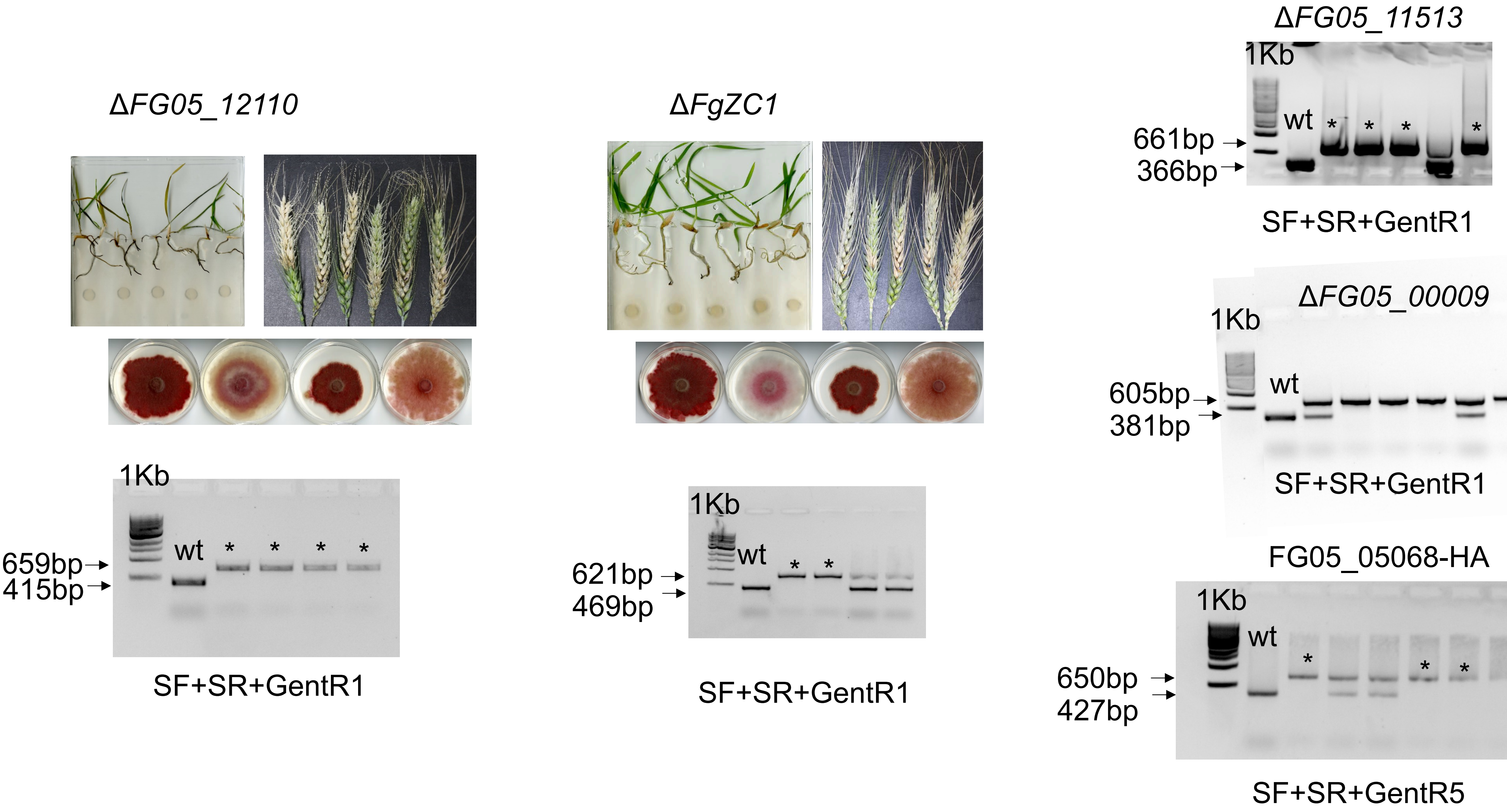
ΔFG05_12110
1Kb
wt
659bp
415bp
SF+SR+GentR1
ΔFgZC1
1Kb
wt
621bp
469bp
SF+SR+GentR1
ΔFG05_11513
1Kb
wt
661bp
366bp
SF+SR+GentR1
ΔFG05_00009
1Kb
wt
605bp
381bp
SF+SR+GentR1
FG05_05068-HA
1Kb
wt
650bp
427bp
SF+SR+GentR5

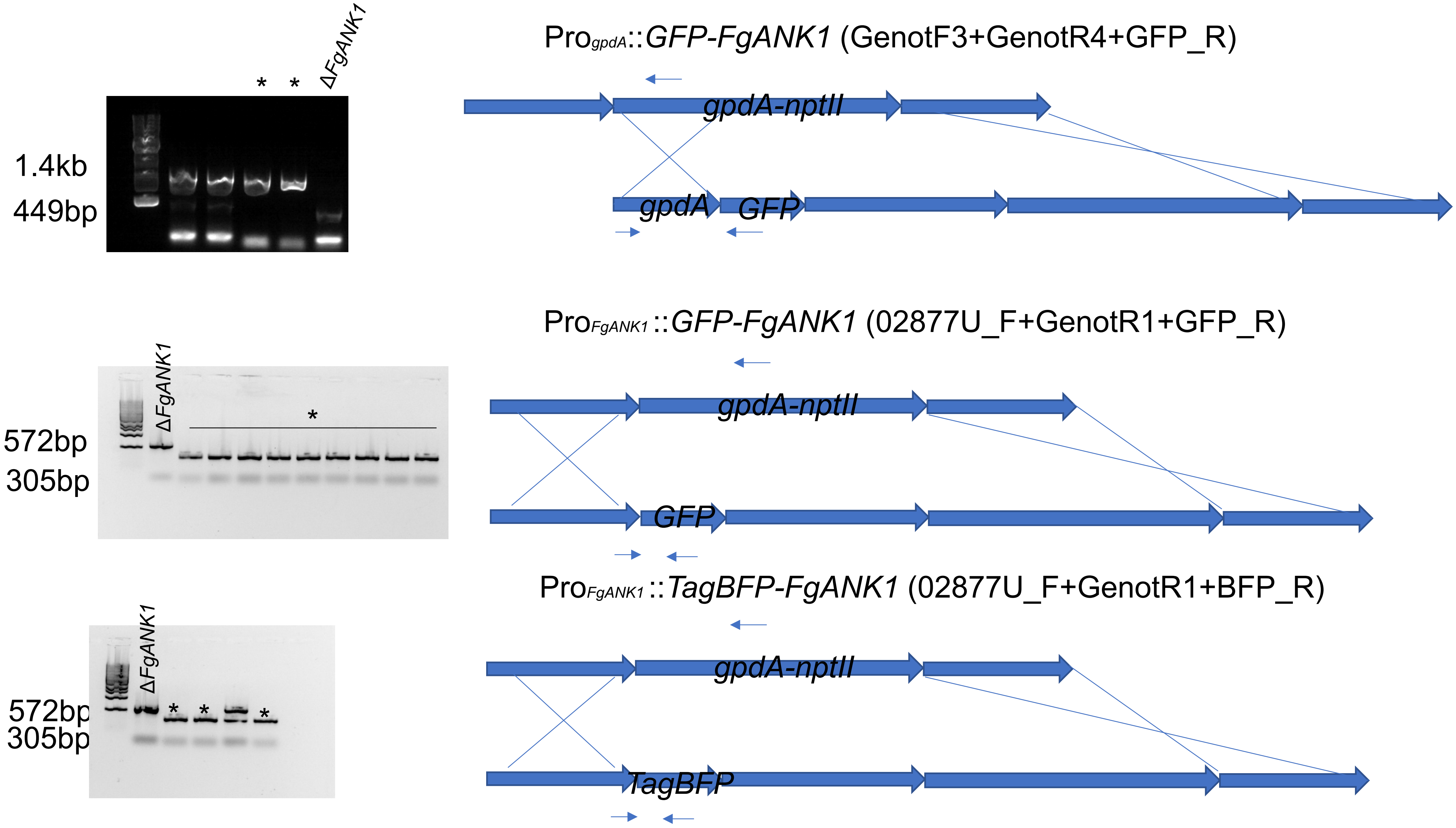

1.4kb
449bp
* *
ΔFgANK1
Pro$_{gpdA}$::GFP-FgANK1 (GenotF3+GenotR4+GFP_R)
gpdA-nptII
gpdA
GFP
572bp
305bp
ΔFgANK1
*
Pro$_{FgANK1}$::GFP-FgANK1 (02877U_F+GenotR1+GFP_R)
gpdA-nptII
GFP
572bp
305bp
ΔFgANK1
* * *
Pro$_{FgANK1}$::TagBFP-FgANK1 (02877U_F+GenotR1+BFP_R)
gpdA-nptII
TagBFP

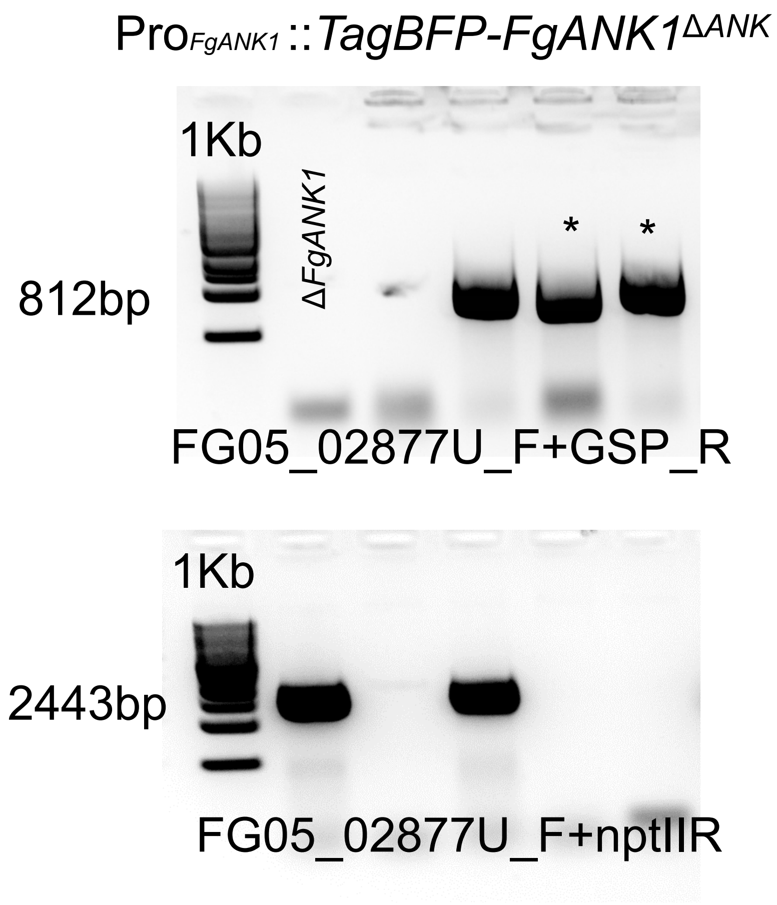
ProFgANK1::TagBFP-FgANK1ΔANK
1Kb
ΔFgANK1
812bp
*
*
FG05_02877U_F+GSP_R
1Kb
2443bp
FG05_02877U_F+nptIIR

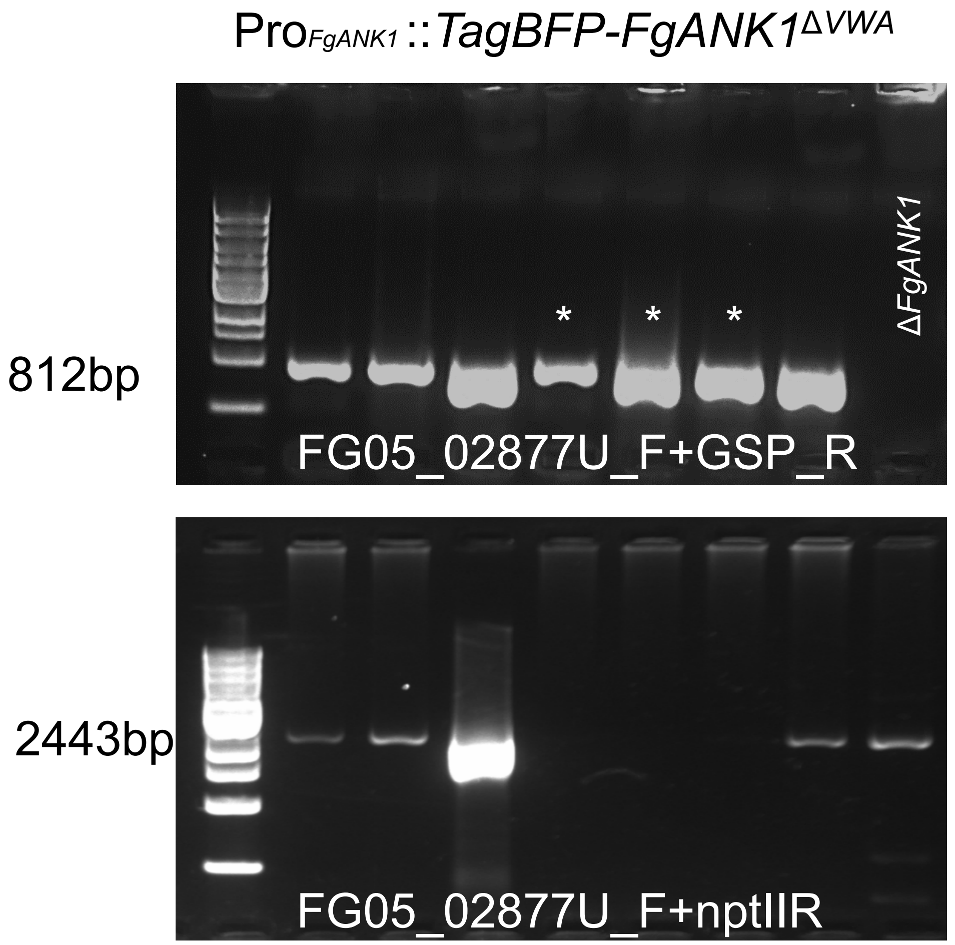
ProFgANK1::TagBFP-FgANK1ΔVWA
812bp
*
*
*
ΔFgANK1
FG05_02877U_F+GSP_R
2443bp
FG05_02877U_F+nptIIR

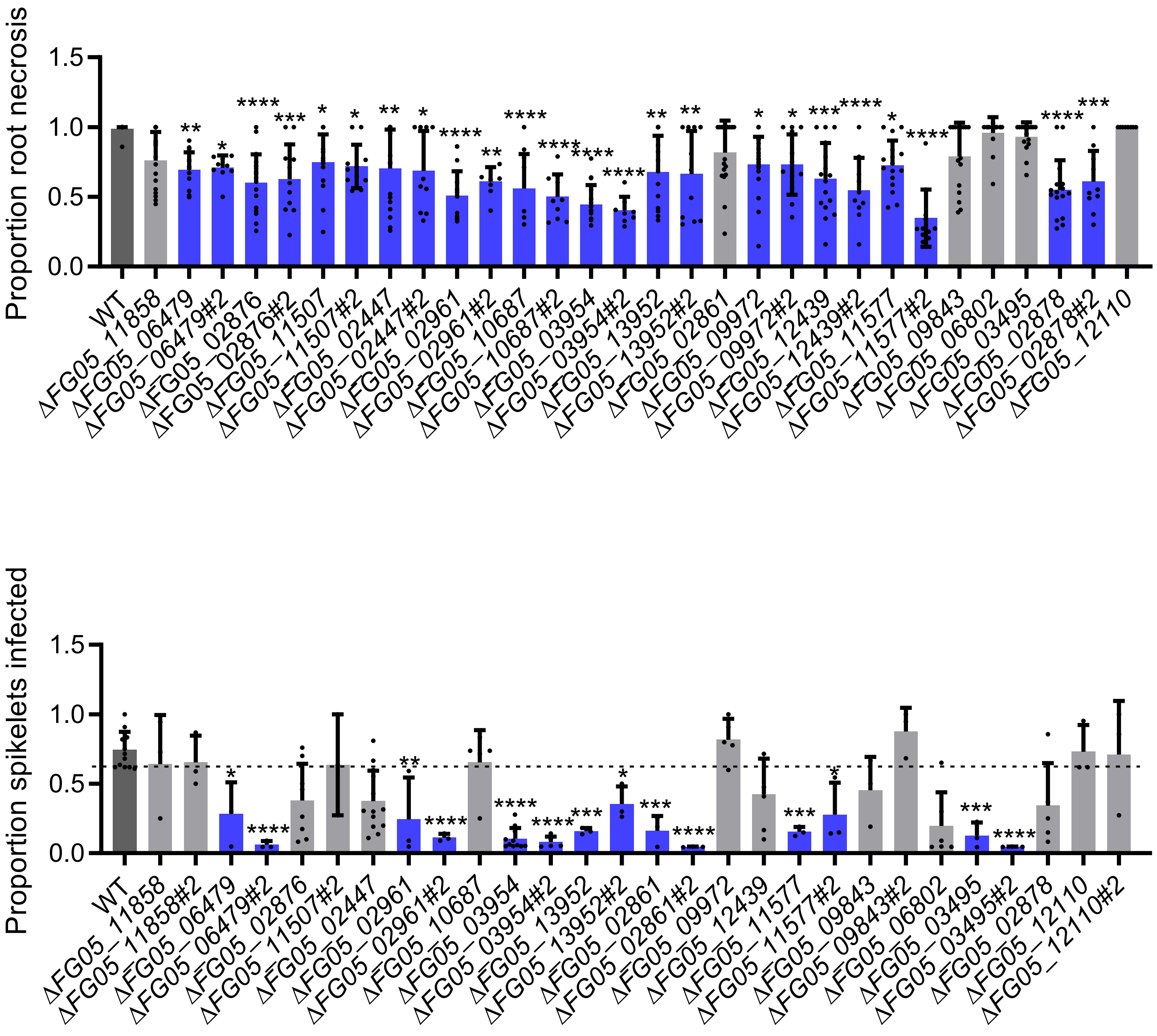
Proportion root necrosis
1.5
1.0
0.5
0.0
WT
ΔFG05_11858
ΔFG05_06479
ΔFG05_06479#2
ΔFG05_02876
ΔFG05_02876#2
ΔFG05_11507
ΔFG05_11507#2
ΔFG05_02447
ΔFG05_02447#2
ΔFG05_02961
ΔFG05_02961#2
ΔFG05_10687
ΔFG05_10687#2
ΔFG05_03954
ΔFG05_03954#2
ΔFG05_13952
ΔFG05_13952#2
ΔFG05_02861
ΔFG05_09972
ΔFG05_09972#2
ΔFG05_12439
ΔFG05_12439#2
ΔFG05_11577
ΔFG05_11577#2
ΔFG05_09843
ΔFG05_06802
ΔFG05_03495
ΔFG05_02878
ΔFG05_02878#2
ΔFG05_12110
Proportion spikelets infected
1.5
1.0
0.5
0.0
WT
ΔFG05_11858
ΔFG05_11858#2
ΔFG05_06479
ΔFG05_06479#2
ΔFG05_02876
ΔFG05_11507#2
ΔFG05_02447
ΔFG05_02961
ΔFG05_02961#2
ΔFG05_10687
ΔFG05_03954
ΔFG05_03954#2
ΔFG05_13952
ΔFG05_13952#2
ΔFG05_02861
ΔFG05_02861#2
ΔFG05_09972
ΔFG05_12439
ΔFG05_11577
ΔFG05_11577#2
ΔFG05_09843
ΔFG05_09843#2
ΔFG05_06802
ΔFG05_03495
ΔFG05_03495#2
ΔFG05_02878
ΔFG05_12110
ΔFG05_12110#2

Suppl. Fig. 5

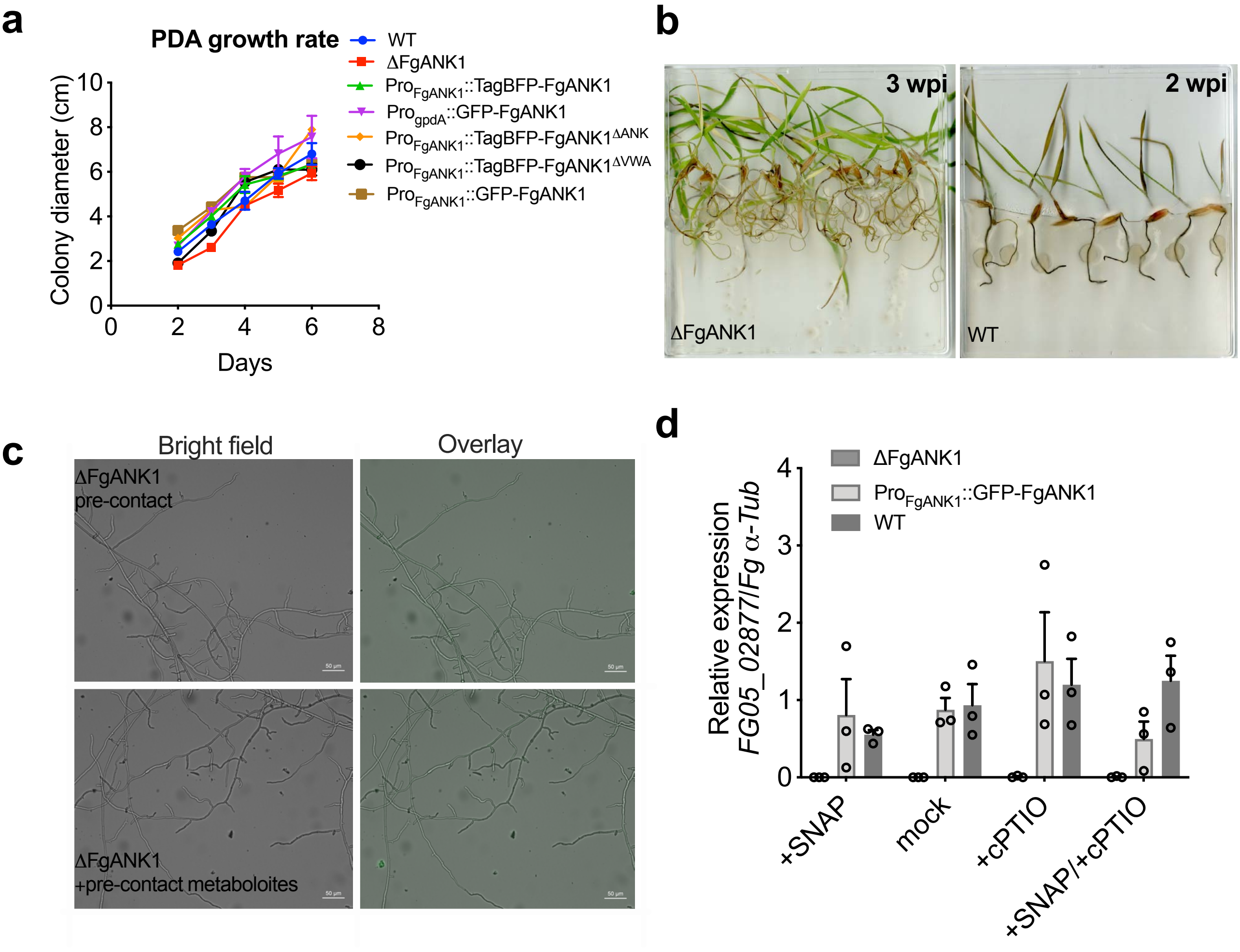
a
PDA growth rate
WT
ΔFgANK1
Pro_FgANK1::TagBFP-FgANK1
Pro_gpdA::GFP-FgANK1
Pro_FgANK1::TagBFP-FgANK1^ΔANK
Pro_FgANK1::TagBFP-FgANK1^ΔVWA
Pro_FgANK1::GFP-FgANK1
Colony diameter (cm)
Days
0
2
4
6
8
10
b
3 wpi
2 wpi
ΔFgANK1
WT
c
Bright field
Overlay
ΔFgANK1
pre-contact
ΔFgANK1
+pre-contact metaboloites
d
ΔFgANK1
Pro_FgANK1::GFP-FgANK1
WT
Relative expression
FG05_02877/Fg α-Tub
0
1
2
3
4
+SNAP
mock
+cPTIO
+SNAP/+cPTIO

### Suppl. Fig. 6

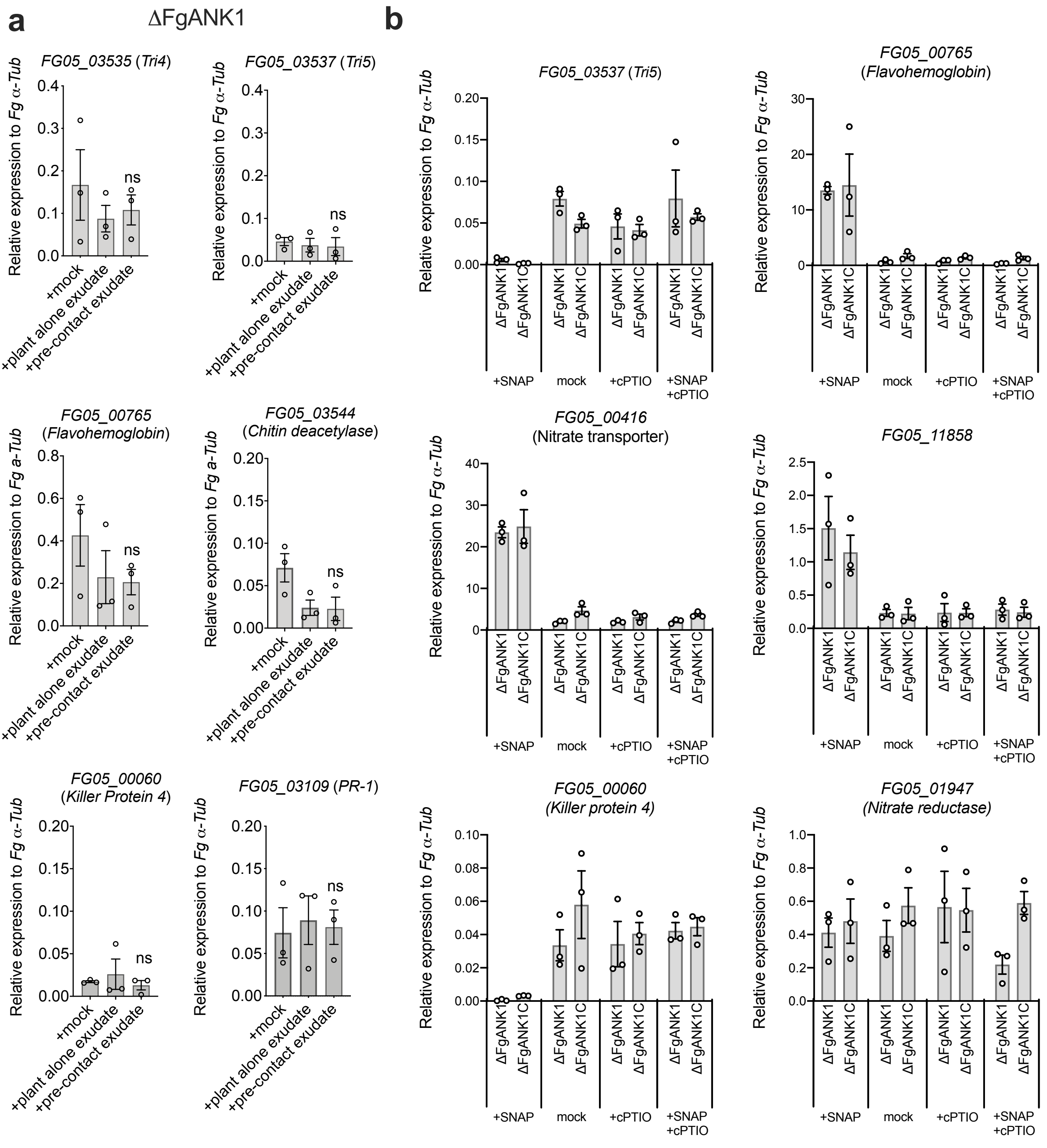
a
ΔFgANK1
FG05_03535 (Tri4)
FG05_03537 (Tri5)
FG05_00765 (Flavohemoglobin)
FG05_03544 (Chitin deacetylase)
FG05_00060 (Killer Protein 4)
FG05_03109 (PR-1)
Relative expression to Fg α-Tub
+mock
+plant alone exudate
+pre-contact exudate
ns
b
FG05_03537 (Tri5)
FG05_00765 (Flavohemoglobin)
FG05_00416 (Nitrate transporter)
FG05_11858
FG05_00060 (Killer protein 4)
FG05_01947 (Nitrate reductase)
Relative expression to Fg α-Tub
ΔFgANK1
ΔFgANK1C
+SNAP
mock
+cPTIO
+SNAP +cPTIO

Suppl. Fig. 7

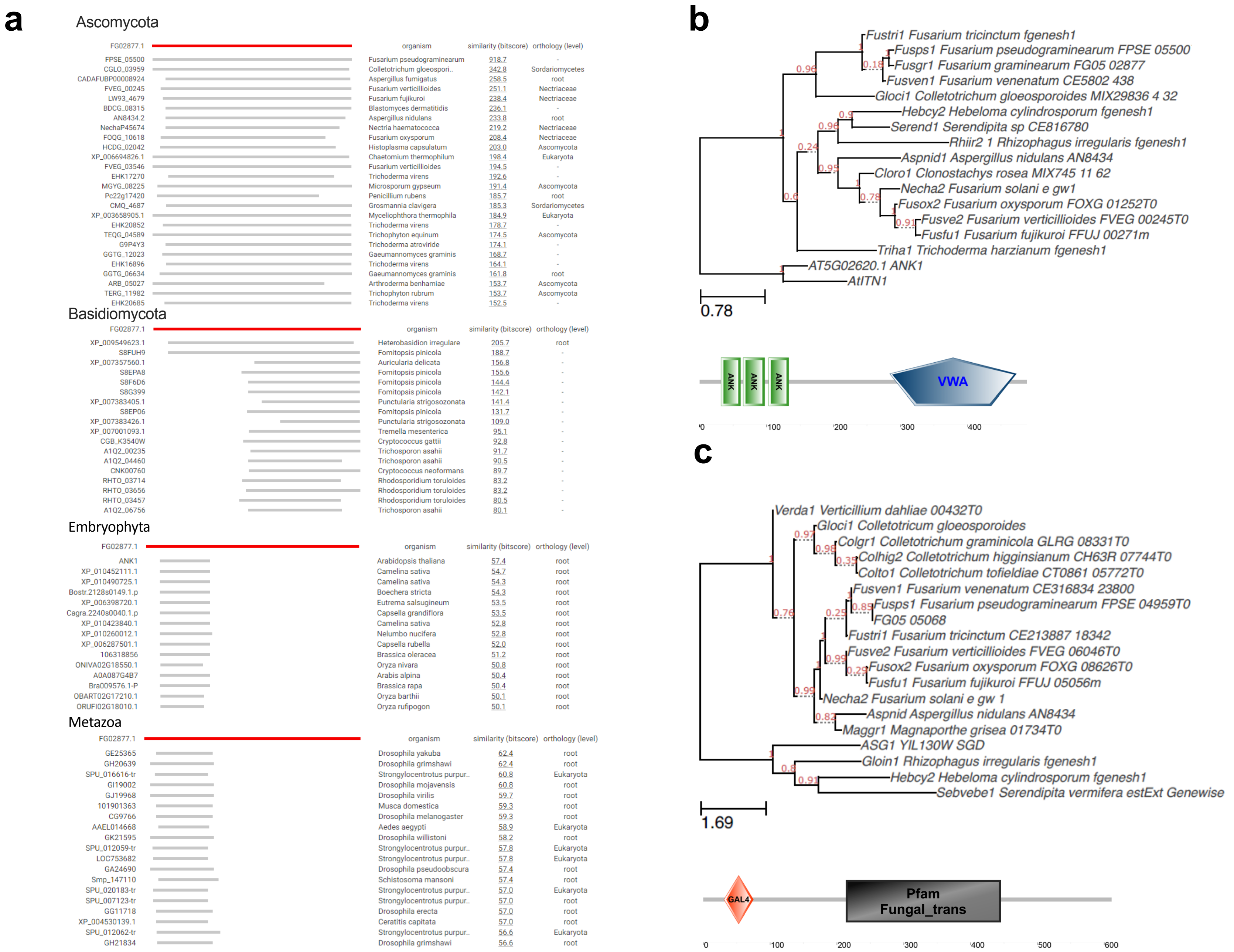
a
Ascomycota
FG02877.1 organism similarity (bitscore) orthology (level)
FPSE_05500 Fusarium pseudograminearum 918.7 -
CGLO_03959 Colletotrichum gloeospori.. 342.8 Sordariomycetes
CADAFUBP00008924 Aspergillus fumigatus 258.5 root
FVEG_00245 Fusarium verticillioides 251.1 Nectriaceae
LW93_4679 Fusarium fujikuroi 238.4 Nectriaceae
BDCG_08315 Blastomyces dermatitidis 236.1 -
AN8434.2 Aspergillus nidulans 233.8 root
NechaP45674 Nectria haematococca 219.2 Nectriaceae
FOQG_10618 Fusarium oxysporum 208.4 Nectriaceae
HCDG_02042 Histoplasma capsulatum 203.0 Ascomycota
XP_006694826.1 Chaetomium thermophilum 198.4 Eukaryota
FVEG_03546 Fusarium verticillioides 194.5 -
EHK17270 Trichoderma virens 192.6 -
MGYG_08225 Microsporum gypseum 191.4 Ascomycota
Pc22g17420 Penicillium rubens 185.7 root
CMQ_4687 Grosmannia clavigera 185.3 Sordariomycetes
XP_003658905.1 Myceliophthora thermophila 184.9 Eukaryota
EHK20852 Trichoderma virens 178.7 -
TEQG_04589 Trichophyton equinum 174.5 Ascomycota
G9P4Y3 Trichoderma atroviride 174.1 -
GGTG_12023 Gaeumannomyces graminis 168.7 -
EHK16896 Trichoderma virens 164.1 -
GGTG_06634 Gaeumannomyces graminis 161.8 root
ARB_05027 Arthroderma benhamiae 153.7 Ascomycota
TERG_11982 Trichophyton rubrum 153.7 Ascomycota
EHK20685 Trichoderma virens 152.5 -
Basidiomycota
FG02877.1 organism similarity (bitscore) orthology (level)
XP_009549623.1 Heterobasidion irregulare 205.7 root
S8FUH9 Fomitopsis pinicola 188.7 -
XP_007357560.1 Auricularia delicata 156.8 -
S8EPA8 Fomitopsis pinicola 155.6 -
S8F6D6 Fomitopsis pinicola 144.4 -
S8G399 Fomitopsis pinicola 142.1 -
XP_007383405.1 Punctularia strigosozonata 141.4 -
S8EP06 Fomitopsis pinicola 131.7 -
XP_007383426.1 Punctularia strigosozonata 109.0 -
XP_007001093.1 Tremella mesenterica 95.1 -
CGB_K3540W Cryptococcus gattii 92.8 -
A1Q2_00235 Trichosporon asahii 91.7 -
A1Q2_04460 Trichosporon asahii 90.5 -
CNK00760 Cryptococcus neoformans 89.7 -
RHTO_03714 Rhodosporidium toruloides 83.2 -
RHTO_03656 Rhodosporidium toruloides 83.2 -
RHTO_03457 Rhodosporidium toruloides 80.5 -
A1Q2_06756 Trichosporon asahii 80.1 -
Embryophyta
FG02877.1 organism similarity (bitscore) orthology (level)
ANK1 Arabidopsis thaliana 57.4 root
XP_010452111.1 Camelina sativa 54.7 root
XP_010490725.1 Camelina sativa 54.3 root
Bostr.2128s0149.1.p Boechera stricta 54.3 root
XP_006398720.1 Eutrema salsugineum 53.5 root
Cagra.2240s0040.1.p Capsella grandiflora 53.5 root
XP_010423840.1 Camelina sativa 52.8 root
XP_010260012.1 Nelumbo nucifera 52.8 root
XP_006287501.1 Capsella rubella 52.0 root
106318856 Brassica oleracea 51.2 root
ONIVA02G18550.1 Oryza nivara 50.8 root
A0A087G4B7 Arabis alpina 50.4 root
Bra009576.1-P Brassica rapa 50.4 root
OBART02G17210.1 Oryza barthii 50.1 root
ORUFI02G18010.1 Oryza rufipogon 50.1 root
Metazoa
FG02877.1 organism similarity (bitscore) orthology (level)
GE25365 Drosophila yakuba 62.4 root
GH20639 Drosophila grimshawi 62.4 root
SPU_016616-tr Strongylocentrotus purpur.. 60.8 Eukaryota
GI19002 Drosophila mojavensis 60.8 root
GJ19968 Drosophila virilis 59.7 root
101901363 Musca domestica 59.3 root
CG9766 Drosophila melanogaster 59.3 root
AAEL014668 Aedes aegypti 58.9 Eukaryota
GK21595 Drosophila willistoni 58.2 root
SPU_012059-tr Strongylocentrotus purpur.. 57.8 Eukaryota
LOC753682 Strongylocentrotus purpur.. 57.8 Eukaryota
GA24690 Drosophila pseudoobscura 57.4 root
Smp_147110 Schistosoma mansoni 57.4 root
SPU_020183-tr Strongylocentrotus purpur.. 57.0 Eukaryota
SPU_007123-tr Strongylocentrotus purpur.. 57.0 root
GG11718 Drosophila erecta 57.0 root
XP_004530139.1 Ceratitis capitata 57.0 root
SPU_012062-tr Strongylocentrotus purpur.. 56.6 Eukaryota
GH21834 Drosophila grimshawi 56.6 root
b
Fustri1 Fusarium tricinctum fgenesh1
Fusps1 Fusarium pseudograminearum FPSE 05500
Fusgr1 Fusarium graminearum FG05 02877
Fusven1 Fusarium venenatum CE5802 438
Gloci1 Colletotrichum gloeosporoides MIX29836 4 32
Hebcy2 Hebeloma cylindrosporum fgenesh1
Serend1 Serendipita sp CE816780
Rhiir2 1 Rhizophagus irregularis fgenesh1
Aspnid1 Aspergillus nidulans AN8434
Cloro1 Clonostachys rosea MIX745 11 62
Necha2 Fusarium solani e gw1
Fusox2 Fusarium oxysporum FOXG 01252T0
Fusve2 Fusarium verticillioides FVEG 00245T0
Fusfu1 Fusarium fujikuroi FFUJ 00271m
Triha1 Trichoderma harzianum fgenesh1
AT5G02620.1 ANK1
AtITN1
0.78
ANK
ANK
ANK
VWA
0
100
200
300
400
c
Verda1 Verticillium dahliae 00432T0
Gloci1 Colletotricum gloeosporoides
Colgr1 Colletotrichum graminicola GLRG 08331T0
Colhig2 Colletotrichum higginsianum CH63R 07744T0
Colto1 Colletotrichum tofieldiae CT0861 05772T0
Fusven1 Fusarium venenatum CE316834 23800
Fusps1 Fusarium pseudograminearum FPSE 04959T0
FG05 05068
Fustri1 Fusarium tricinctum CE213887 18342
Fusve2 Fusarium verticillioides FVEG 06046T0
Fusox2 Fusarium oxysporum FOXG 08626T0
Fusfu1 Fusarium fujikuroi FFUJ 05056m
Necha2 Fusarium solani e gw 1
Aspnid Aspergillus nidulans AN8434
Maggr1 Magnaporthe grisea 01734T0
ASG1 YIL130W SGD
Gloin1 Rhizophagus irregularis fgenesh1
Hebcy2 Hebeloma cylindrosporum fgenesh1
Sebvebe1 Serendipita vermifera estExt Genewise
1.69
GAL4
Pfam
Fungal_trans
0
100
200
300
400
500
600

Suppl. Fig. 8

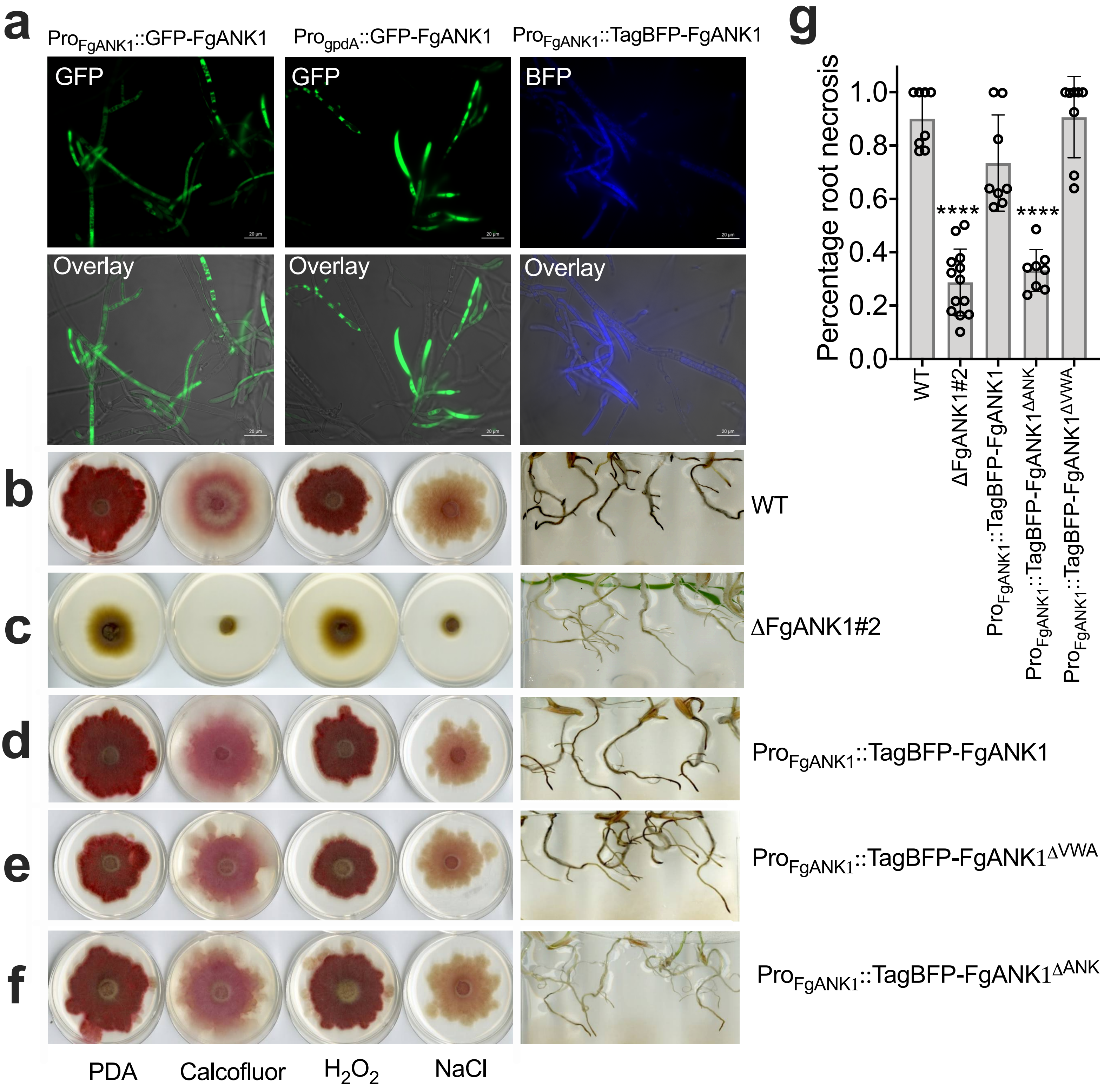

a
b
c
d
e
f
g
Pro$_{FgANK1}$::GFP-FgANK1
Pro$_{gpdA}$::GFP-FgANK1
Pro$_{FgANK1}$::TagBFP-FgANK1
GFP
GFP
BFP
Overlay
Overlay
Overlay
WT
ΔFgANK1#2
Pro$_{FgANK1}$::TagBFP-FgANK1
Pro$_{FgANK1}$::TagBFP-FgANK1$^{ΔVWA}$
Pro$_{FgANK1}$::TagBFP-FgANK1$^{ΔANK}$
PDA
Calcofluor
$H_2O_2$
NaCl
Percentage root necrosis
1.0
0.8
0.6
0.4
0.2
0.0
****
****
WT
ΔFgANK1#2
Pro$_{FgANK1}$::TagBFP-FgANK1
Pro$_{FgANK1}$::TagBFP-FgANK1$^{ΔANK}$
Pro$_{FgANK1}$::TagBFP-FgANK1$^{ΔVWA}$

### Suppl. Fig. 9

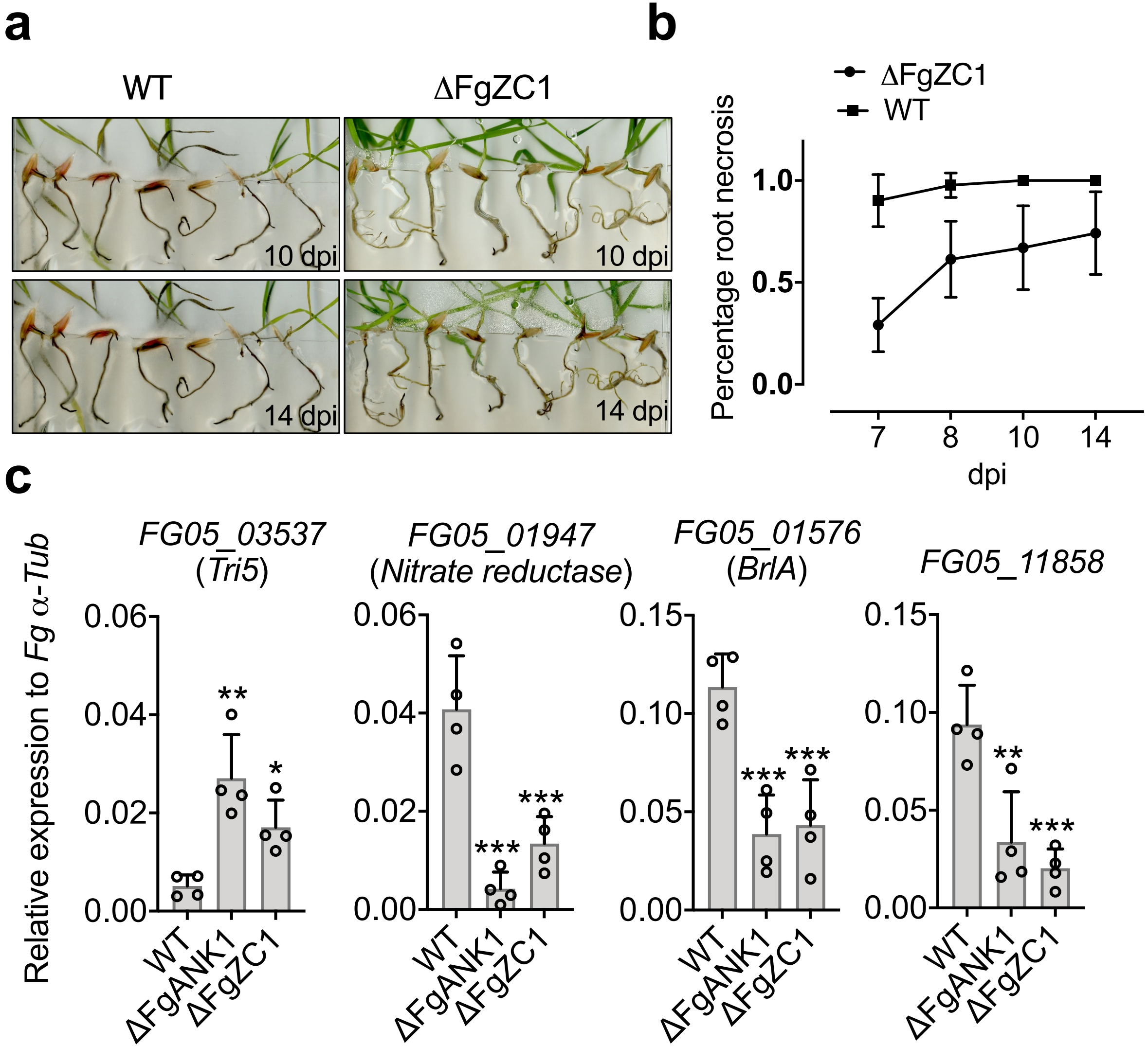
a
WT
ΔFgZC1
10 dpi
10 dpi
14 dpi
14 dpi
b
ΔFgZC1
WT
Percentage root necrosis
1.0
0.5
0.0
7
8
10
14
dpi
c
Relative expression to Fg α-Tub
FG05_03537
(Tri5)
FG05_01947
(Nitrate reductase)
FG05_01576
(BrlA)
FG05_11858
0.06
0.04
0.02
0.00
0.15
0.10
0.05
0.00
**
*
***
***
***
***
**
***
WT
ΔFgANK1
ΔFgZC1
